# PRMT1-SFPQ regulates intron retention to control matrix gene expression during craniofacial development

**DOI:** 10.1101/2024.07.09.598733

**Authors:** Julia Raulino Lima, Nicha Ungvijanpunya, Qing Chen, Hoang Quoc Hai Pham, Tal Rosen, Greg Park, Mohammadreza Vatankhah, Steven Yen, Yang Chai, Amy E. Merrill, Zhaoyang Liu, Jian-Fu Chen, Yanzhong Yang, Weiqun Peng, Jian Xu

## Abstract

Spliceosomopathies, which are a group of disorders caused by defects in the splicing machinery, frequently affect the craniofacial skeleton and limb, but the molecular mechanism underlying this tissue-specific sensitivity remains unclear. Splicing factors and small nuclear ribonucleoproteins (snRNPs) are core components of splicing machinery, and splicing factors are further controlled by post-translational modifications, among which arginine methylation is one of the most prevalent. We determined the splicing mechanisms in the cranial neural crest cells (CNCCs), a multipotent developmental population that gives rise to the majority of the craniofacial skeleton, and focused on an upstream regulator of splicing proteins, protein arginine methyltransferase 1 (PRMT1). PRMT1 is the highest expressing arginine methyltransferase in CNCCs and its role in craniofacial development is evident from our earlier investigation, where CNCC-specific *Prmt1* deletion caused cleft palate and mandibular hypoplasia. PRMT1 catalyzes arginine methylation of splicing factors to modify protein localization, expression and activity. In the present study, we uncover roles of PRMT1 in the regulation of intron retention, a type of alternative splicing where introns are retained in the mature mRNA. CNCCs from mandibular primordium of *Prmt1*-deficient embryos demonstrated an increase in the percentage of intron-retaining mRNA of matrix genes, which triggered NMD, causing a reduction in matrix mRNA abundance. We further identified SFPQ as a substrate of PRMT1 that depends on PRMT1 for arginine methylation and protein expression in the developing craniofacial structures. Depletion of SFPQ in CNCCs phenocopied PRMT1 deletion whereby matrix, Wnt signaling components and neuronal gene transcripts contained higher IR and exhibited lower expression. We further recognized gene length as a common feature among SFPQ-regulated genes in CNCCs. Altogether, these findings demonstrate that the PRMT1-SFPQ pathway modulates matrix gene expression via IR-triggered NMD in CNCCs during craniofacial development.

## INTRODUCTION

Craniofacial abnormalities affecting bone formation in the skull and face are the most common birth defects in infants. Proper formation of these structures involves coordination of proliferation, migration, and differentiation of cranial neural crest cells (CNCCs) (Martik & Bronner, 2017; Plein, Fantin, & Ruhrberg, 2015). CNCCs are a transient population of progenitor cells that populate the first and second pharyngeal arches and give rise to the facial skeleton including maxilla, mandible and palates, and the anterior skull (Chai et al., 2000; Jiang, Iseki, Maxson, Sucov, & Morriss-Kay, 2002; Martik & Bronner, 2017). Dysregulation of transcription factors, chromatin remodelers, RNA regulatory proteins, and signaling molecules have been implicated in impaired neural crest development that results in craniofacial defects (Bélanger et al., 2018; Martik & Bronner, 2017; Plein et al., 2015; Strobl-Mazzulla, Marini, & Buzzi, 2012). Spliceosomopathies, which are a group of disorders caused by defects in the splicing machinery, frequently affect the craniofacial skeleton and limb. However, the mechanisms behind the sensitivity of these tissues to spliceosomal defects remain incompletely understood. Splicing factors and small nuclear ribonucleoproteins (snRNPs) are core components of splicing machinery, which control pre-mRNA splicing and mRNA maturation through the process of alternative splicing (AS). AS includes seven types of events, exon skipping, mutually exclusive exons, alternative 5’ and 3’-splice sites, alternative promoters, alternative polyadenylation, and intron retention (Hooper, Jones, Smith, Williams, & Li, 2020). Intron retention (IR), where introns are retained within the mature mRNA sequence, is abundant in plants and fungi (E. T. Wang et al., 2008). Although initially overlooked in mammals due to challenges in analysis, recent technical advances have unveiled its significant roles in various aspects, including cell differentiation and stress response, especially in neuronal, immune, and cancer cells (Braunschweig et al., 2014; Marquez, Brown, Simpson, Barta, & Kalyna, 2012; Monteuuis, Wong, Bailey, Schmitz, & Rasko, 2019; Wong & Schmitz, 2022).

In exploring the splicing mechanisms in CNCCs, we focused on upstream regulators for splicing factors. The activity of splicing factors is tightly controlled by post-translational modifications, among which methylation is the most abundant, surpassing phosphorylation (Ruta, Pagliarini, & Sette, 2021; Thandapani, O’Connor, Bailey, & Richard, 2013). Splicing regulator methylation mostly occurs on arginine motifs, catalyzed by the protein arginine methyltransferase (PRMT) family of enzymes. These methylation events determine their protein stability, subcellular localization, and splicing activity, shaping the splicing product (Blanc & Richard, 2017; Liu & Dreyfuss, 1995; Snijders et al., 2015). PRMT1 is the most abundant PRMT in most cell types. It specifically catalyzes methylation of arginine (R) residues within RG/RGG/GAR repeats, which are motifs enriched in RNA-binding proteins, particularly splicing regulators. This results in asymmetric dimethylation of arginine (ADMA) (Smith, Schurter, Wong-Staal, & David, 2004; Thandapani et al., 2013). The importance of PRMT1 in craniofacial development is evident from our earlier investigation in which genetic deletion of *Prmt1* in CNCCs leads to cleft palate and mandibular hypoplasia (Gou, Li, Jackson-Weaver, et al., 2018; Gou, Li, Wu, et al., 2018). In the present study, we uncovered a previously unrecognized role of PRMT1 in regulating intron retention in CNCCs. The mandibular primordium of *Prmt1*-deficient embryos demonstrated increased retention of introns in mature mRNAs that encode bone and cartilage matrix components. These retained introns contain premature termination codons (PTCs) that trigger nonsense-mediate decay (NMD) to degrade mRNA and reduce gene expression. We further identified SFPQ, EWSR1 and TAF15 as downstream substrates of PRMT1 in the developing craniofacial structures, which depends on PRMT1 for arginine methylation. SFPQ further depends on PRMT1 for protein stability. Depletion of SFPQ in CNCCs partially phenocopied PRMT1 deletion, causing elevated retention of introns in mRNA of matrix, neuronal, and Wnt signaling pathway genes. Matrix and Wnt pathway genes susceptible to perturbation of the PRMT1-SFPQ pathway are further characterized as long genes with a median length of 100kb. Their retained introns trigger NMD to reduce gene expression. Together, these findings demonstrate that the PRMT1-SFPQ pathway modulates CNCC gene expression via control of splicing. Deficiency of this pathway leads to aberrant intron retention, which induces mRNA decay that downregulates matrix expression in CNCCs during craniofacial development.

## RESULTS

### CNCCs from the embryonic mandibular process display abundant intron retention (IR), which was further elevated by loss of *Prmt1*

PRMT1 is the most abundant enzyme of the arginine methyltransferase family in cranial neural crest cells (CNCCs), with an expression level ∼10-100 folds higher than other PRMTs (Figure-1A). We previously demonstrated critical roles for PRMT1 in CNCCs during embryogenesis using genetic deletion, whereby neural crest-specific deletion of *Prmt1* caused cleft palate and shorter mandibles in mouse (Gou, Li, Jackson-Weaver, et al., 2018; Gou, Li, Wu, et al., 2018). Shorter mandibles, or mandibular hypoplasia is a phenotype shared by most spliceosomopathies, a group of conditions caused by splicing defects (Griffin & Saint-Jeannet, 2020; Merkuri & Fish, 2019). To investigate the molecular mechanisms underlying PRMT1’s role in mandible development and splicing regulation, we conducted a genome-wide analysis of transcriptional and splicing alterations within the embryonic mandibles following *Prmt1* deletion. For this purpose, CNCCs, the cell lineage that gives rise to bone and cartilage in the mandible, were isolated from mandibular processes of control (*Wnt1-Cre; R26R ^tdTomato^*) or *Prmt1* deficient (*Prmt1 CKO, Wnt1-Cre; Prmt1^fl/fl^; R26R ^tdTomato^*) embryos at embryonic day 13.5 (E13.5) for mRNA extraction and sequencing (Figure-1B, Supplemental Figure-S1). The ensuing comparative analysis using rMATS (Y. Wang et al., 2024) unveiled changes across five types of alternative splicing events (Figure-1C), echoing our earlier finding where PRMT1 regulates exon usage in epicardial cells during cardiac morphogenesis (Jackson-Weaver et al., 2020). In contrast to embryonic epicardial cells, we noted a higher abundance of IR changes in mandibular CNCCs (Figure-1C). The IR changes were exemplified by the increase of intron retention in *Pex12, Mmp23,* and *Ecm1* (Figure-1D, 1F), and the decrease of retained intronic expression in *Tbx1* (Figure-1E, 1F). To comprehensively analyze intron retention at a gene-specific level, we employed IRI (intron reads index), which is an algorithm that quantifies the ratio of normalized read counts between intronic and exonic regions (Ni et al., 2016; Sun et al., 2023; Tian et al., 2020). IRI analysis demonstrated that *Prmt1* deficiency didn’t cause a global shift in intron retention levels (Supplemental Figure-S2) but changed intron retention in genes of multiple biological pathways (Supplemental Table S1). We noted protein binding, metal ion binding, and cartilage development among genes that present increased intron retention, while mitochondrion and metabolic pathways were identified among genes with decreased intron retention (Supplemental Table S1). These findings identify intron retention as a prevalent phenomenon in CNCCs during craniofacial development, and the deletion of *Prmt1* further enhanced IR in genes involved in diverse biological processes.

**Figure 1.**
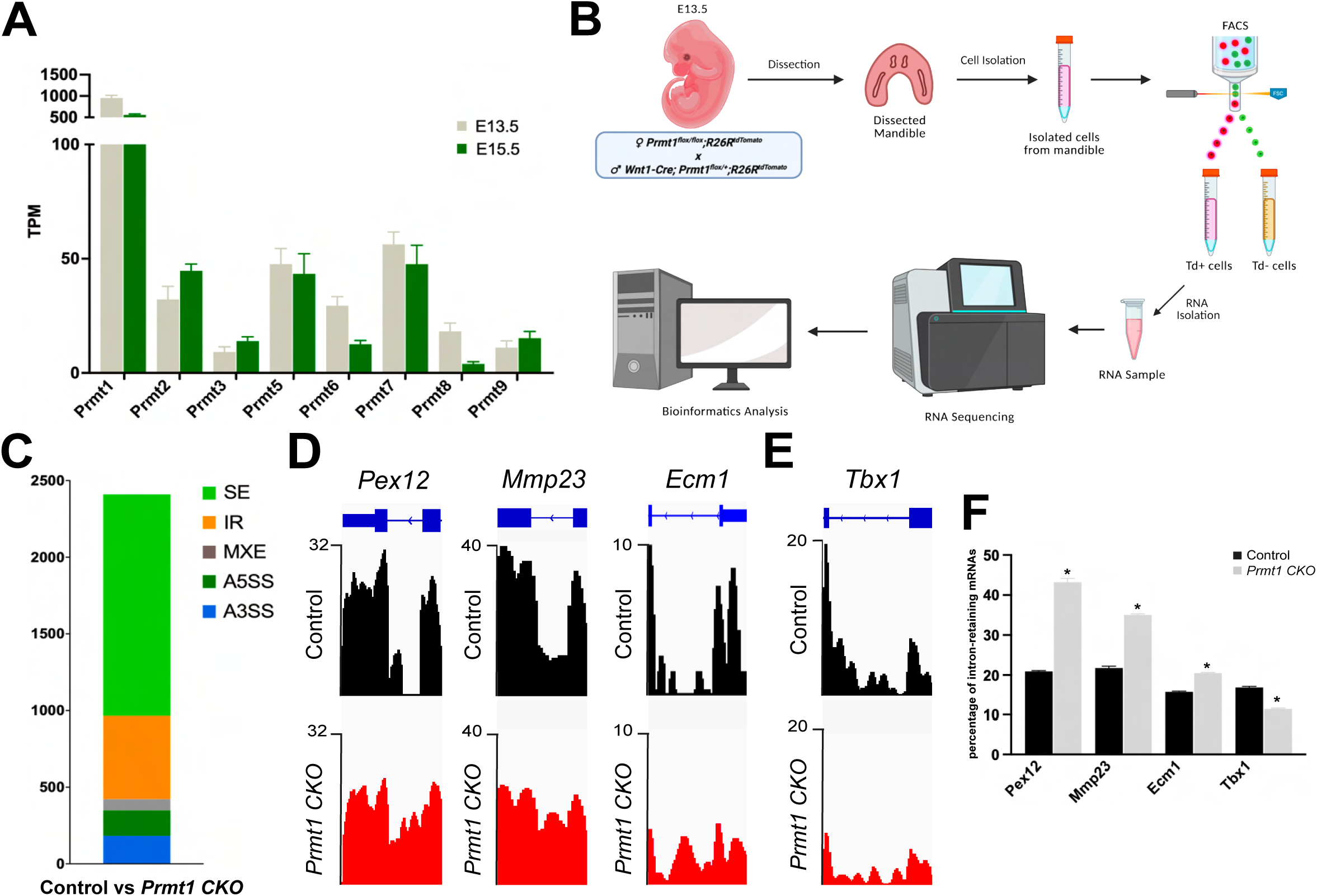
CNCC-specific deletion of *Prmt1* elevates intron retention in the embryonic mandibular process. (A) Expression levels of PRMT1-9 mRNAs in primary isolated cranial neural crest cells (CNCC) from *Wnt1-Cre; R26R^tdTomato^* mouse embryo heads at E13.5 and E15.5. TPM, transcript per million. (B) Diagram illustrating the isolation of CNCCs from embryonic mandibles, followed by poly(A)+ mRNA isolation and sequencing (n=4 in control and *Prmt1 CKO* group). (C) *Prmt1* deletion in CNCC caused changes in alternative splicing (AS). Changes in AS events were analyzed by rMATS using RNA sequencing data and significant changes in each type of AS were shown in a stacked bar chart. SE, skipped exon. IR, intron retention. MXE, mutually exclusive exons. A5SS, alternative 5’ splice site. A3SS, alternative 3’ splice site. (D & E) Intron retention was prevalent in CNCCs and altered by *Prmt1* deletion. Track view of genes demonstrating intronic and exonic expression with blue boxes indicating exons and blue lines indicating introns. Intron retention was elevated by *Prmt1* deletion in *Pex12, Mmp23,* and *Ecm1 (D)* and reduced in *Tbx1 (E)*. (F) Quantification of intron expression in *Pex12, Mmp23, Ecm1* and *Tbx1* by the percentage of intron-retaining mRNAs in each gene, calculated from IRI analysis based on RNA-seq data. * p<0.05 *Prmt1* CKO vs Control. Control: *Wnt1-Cre; R26R^tdTomato^*. *Prmt1 CKO: Wnt1-Cre; Prmt1^fl/fl^; _R26RtdTomato_*.

### Neural crest-specific deletion of *Prmt1* caused a significant reduction of matrix gene expression in the developing mandibles

The main documented role for intron retention is to reduce expression of the gene harboring retained introns via non-sense mediated decay (NMD), the cytosolic RNA surveillance system (Schmitz et al., 2017). When intron-retaining mRNA transcripts are loaded onto the translation machinery, introns that contain premature termination codons (PTCs) within their open reading frames trigger NMD, leading to mRNA transcript degradation. Reduction in mRNA abundance leads to decreased protein expression for genes harboring these introns (Wong, Au, Ritchie, & Rasko, 2016). The intron-retaining transcripts illustrated in Figure-1, *Pex12, Mmp23,* and *Ecm1* demonstrated increased intronic expression and decreased exonic expression, supporting this negative correlation (Figure-1D, 1F). To systematically analyze genes that are differentially downregulated, we compared the transcription profile between control and *Prmt1* CKO mandibular CNCCs and revealed downregulation of 303 genes and concurrent upregulation of 160 genes by *Prmt1* deletion (Figure-2A, 2B). Upon pathway analysis, glycosaminoglycan (GAG) degradation (35 out of 303 genes) and extracellular matrix (ECM) (33 out of 303 genes) emerged as top pathways among downregulated genes (Figure-2C). These GAG degradation and ECM genes encode matrix proteins pivotal to osteogenic/chondrogenic differentiation and the formation of bone and cartilage matrix. Ingenuity Pathway Analysis (IPA) of downregulated genes further suggested a disruption in bone, cartilage, and connective tissue development within *Prmt1* CKO mutant mandibles (Figure-2D). In addition, *Prmt1* deficiency upregulated genes in cytokine-mediated signaling pathway and p53 signaling, which are consistent with previous reports that *Prmt1* deletion resulted in cytokine production in oral epithelium and p53 accumulation in embryonic epicardium (Jackson-Weaver et al., 2020; Zhang et al., 2018). *Prmt1* deletion also upregulated genes involved in adult behavior, postsynaptic membrane organization and regulated exocytosis, which echoes findings on PRMT1 function in the neuronal lineages and cancer (Chen et al., 2021; Hashimoto, Fukamizu, Nakagawa, & Kizuka, 2021; Hashimoto, Kumabe, et al., 2021; Hashimoto et al., 2022; Xu & Richard, 2021) (Figure-2E).

**Figure 2.**
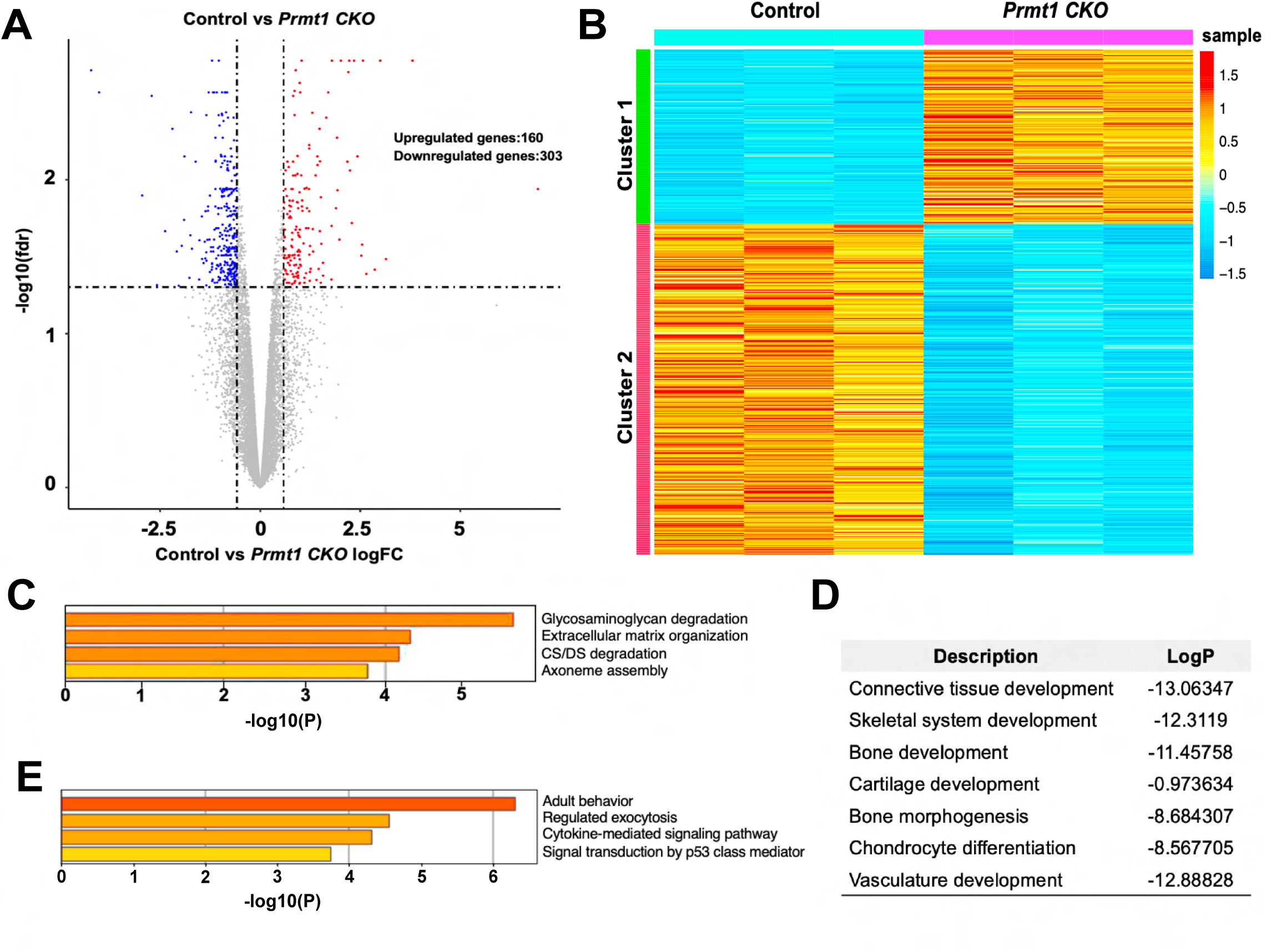
CNCC-specific *Prmt1* deletion reduced matrix gene expression in the developing mandibles. (A) Volcano plot illustrating upregulation of 160 and downregulation of 303 genes in the mandibular primordium of *Prmt1*-deficient embryos at E13.5, compared to control mandibles. (B) Heatmap showing differential gene expression between control and *Prmt1-*deficient mandibles. (C) GO analysis of pathway enrichment in downregulated genes demonstrating glycosaminoglycan (GAG) degradation and extracellular matrix (ECM) organization as the top pathways. (D) Ingenuity Pathway Analysis suggesting connective tissue, bone, and cartilage development as affected biological processes based on downregulated genes. (E) GO analysis of pathway enrichment in upregulated genes demonstrating adult behavior and cytokine-mediated signaling and p53 signal transduction as top pathways. Control: *Wnt1-Cre; R26R^tdTomato^* (n=4). *Prmt1 CKO: Wnt1-Cre; Prmt1^fl/fl^; R26R^tdTomato^* (n=4).

### Deletion of *Prmt1* in CNCCs elevated intron retention in mRNAs that encode ECM and GAG degradation enzymes

To determine whether downregulation of matrix genes in *Prmt1 CKO* embryos is regulated by IR-mediated mechanism, we first focused on ECM genes downregulated in *Prmt1*-deficient mandibles and investigated their intron retention events. The findings are presented in a scatter plot where the red line denotes unchanged IR. A prominent elevation of IR within ECM genes was revealed by the fact that most ECM data points landed above the red line (Figure-3A). ECM genes with the most significant IR increase were labeled and highlighted in red. The IR increase was also exemplified by *Adamts16* and *Cthrc1* (Figure-3B, 3C), where track view of the two genes illustrated increased IR and reduced exonic expression (Figure-3Ba, 3Bb). To validate the reproducibility of IR elevation in ECM transcripts, we collected four additional embryos from each group, isolated mature mRNA using poly(A)+ magnetic beads, and designed primers for intronic and exonic region of additional ECM genes with RT-PCR analysis. In line with our initial observations, intronic expression increased significantly, coupled with decreased exonic expression in *Dcn, Tnn, Lox, Loxl1, Matn2, Col14a1, Adamts12, Adamts2, Adam12,* and *Scara5* of the *Prmt1* CKO CNCCs as compared to control (Figure-3D). The decrease in exonic expression, representing mRNA abundance, led to a decline in protein expression, as demonstrated by immunostaining for LOXL1 and FBLN5 in CNCCs (Figure-3E-3H). These findings demonstrate that ECM transcripts in *Prmt1* CKO mandibles exhibited IR elevation and reduction in mRNA expression, leading to lower ECM protein production. Next, we analyzed the intron retention events in GAG degradation genes, which is the top downregulated gene cluster in the *Prmt1* CKO group and clinically associated with craniofacial birth defects (Mizumoto & Yamada, 2021; Paganini, Gramegna Tota, Superti-Furga, & Rossi, 2020; Schwartz & Domowicz, 2002). Similar to ECM genes, GAG degradation genes showed IR elevation in scatter plot where most data points landed above the red line (Figure-3I). The track view of *St6galnac3* and *Galn11* further illustrated the increased IR and lower mRNA abundance (Figure-3J, 3K). Taken together, ECM and GAG degradation genes that are downregulated in *Prmt1 CKO* embryos demonstrate elevated intron retention.

**Figure 3.**
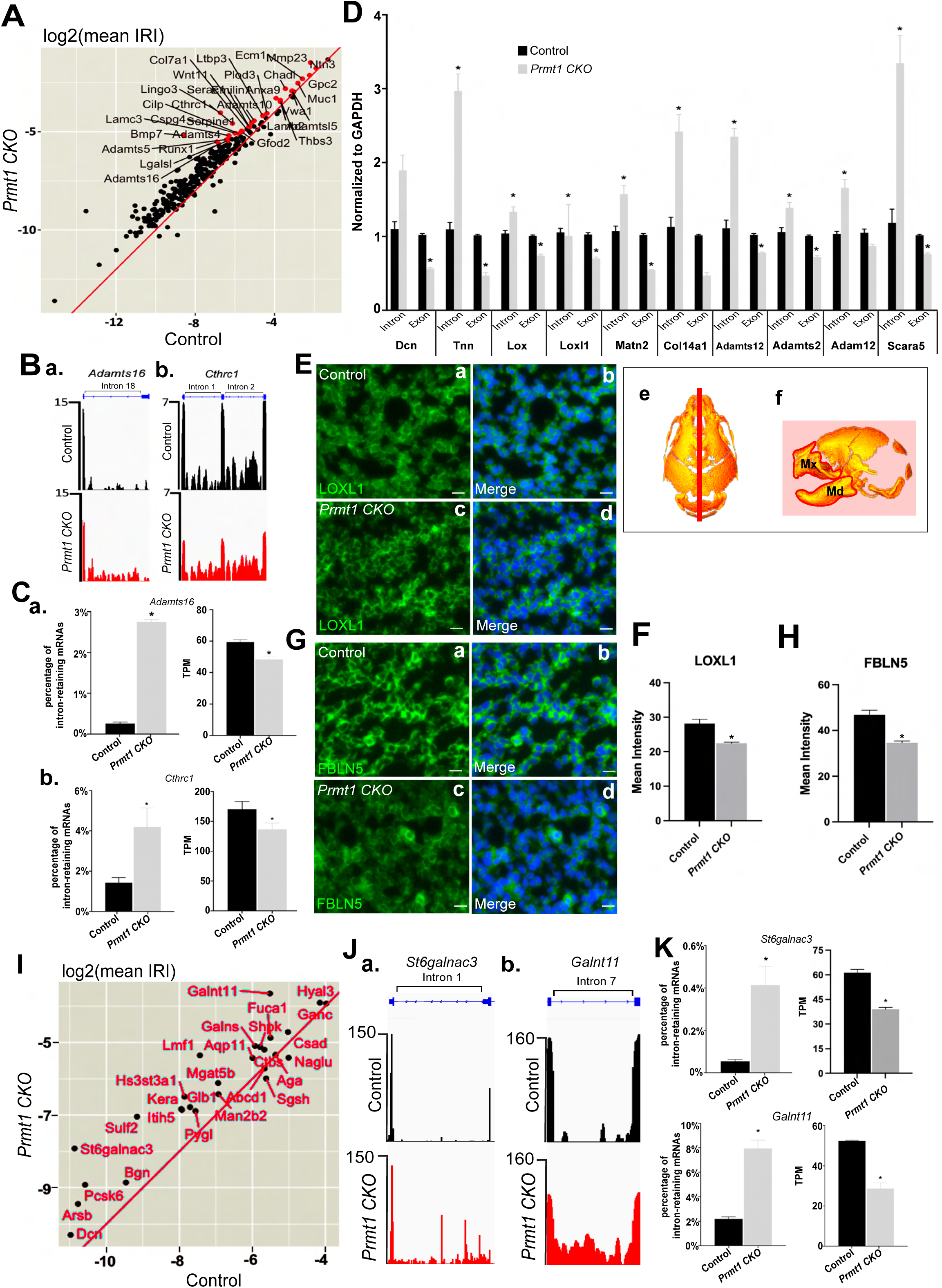
PRMT1 regulates intron retention in ECM and GAG degradation genes. (A-C) Intron retention increased in the majority of ECM gene transcripts that were downregulated in *Prmt1* deficient embryos, as illustrated by a scatter plot based on intron retention index (IRI). The red line delineates unchanged levels of intron retention. Genes with the top differential IR were represented by red dots and labeled. ECM gene transcripts *Adamts16* and *Cthrc1* demonstrating higher intron retention in *Prmt1-* deleted embryos were illustrated by track view in *B* and quantified for intronic (left) and exonic (right) expression as shown in *C*. (D) Higher intron retention and lower mRNA abundance of ECM genes were validated in additional embryo samples by RT-PCR. Primers that span the intronic or intron-exon junction region were used to assess intronic expression. Primers that span the exonic region were used to examine exonic expression that indicated mRNA abundance. (E-H) Reduced expression of LOXL1 and FBLN5 was examined at the protein level by immunostaining (*E, G*) and quantified (*F, H*). Ee and Ef illustrated the plane of section and the region of analysis for *E* and *G*. (I-K) Intron retention increased in the majority of GAG degradation gene transcripts that were downregulated in *Prmt1* deficiency, as indicated by a scatter plot based on IRI. The red line defines where intron retention is unchanged. Genes were represented by black dots and labeled in red. GAG degradation genes *St6galnac3* and *Galnt11* demonstrating higher intron retention in *Prmt1-*deleted embryos were illustrated by track view (*J*) and quantified for intronic (left) and exonic (right) expression (*K*). * p<0.05 *Prmt1* CKO vs Control. Control: *Wnt1-Cre; R26R^tdTomato^*. *Prmt1 CKO: Wnt1-Cre; Prmt1^fl/fl^; R26R^tdTomato^*. Scale bar: (E,G), 100µm.

### IR-triggered NMD acts as a physiological and stress-induced mechanism for mRNA decay in CNCCs

We inspected the retained introns in these ECM and GAG degradation transcripts and identified PTCs within their open reading frames, suggesting PTC-triggered NMD as a potential mechanism for transcript decay (Supplemental Table S2) (Wong et al., 2016). To test this hypothesis, we inhibited the NMD machinery using a chemical inhibitor NMDI14, which disrupts the interaction between UPF1 and SMG7 to block NMD complex formation and activity (Martin et al., 2014). To this end, *Wnt1-Cre; Prmt1^fl/+^; R26R ^tdTomato^* were bred with *Prmt1^fl/fl^; R26R ^tdTomato^* mice to generate *Prmt1 CKO* (*Wnt1-Cre; Prmt1^fl/fl^; R26R ^tdTomato^*) and *Prmt1 heterozygous* embryos as littermate control *(Prmt1 Het, Wnt1-Cre; Prmt1^fl/+^; R26R ^tdTomato^*). Primary CNCCs from E13.5 *Prmt1 CKO* and *Prmt1 Het* were plated and treated with NMDI14 or DMSO as control, followed by poly(A)+ mRNA extraction and RT-PCR analysis using primers that span the intronic or exonic regions. We first validated the inhibition of NMD activity by NMDI14 through analysis of *glutathione peroxidase 1 (Gpx1) intron 1,* which is a well-documented substrate for NMD-mediated transcript decay (Sun, Moriarty, & Maquat, 2000). We further demonstrated that NMDI14 caused a significant increase of *Adamts2* intron 19 and *Alpl* intron 5 expression in the control CNCCs from *Prmt1 Het* (Figure-4Aa), suggesting that inhibition of NMD blocked IR-triggered mRNA decay, thereby causing accumulation of introns. A more prominent accumulation of introns was induced by NMDI14 in the *Prmt1 CKO* CNCCs, accompanied by significant increase of mRNA abundance indicated by higher exonic expression levels (Figure-4Ab). These data suggest that *Prmt1* deficiency invoked IR-induced NMD for mRNA decay which reduced mRNA abundance.

**Figure 4.**
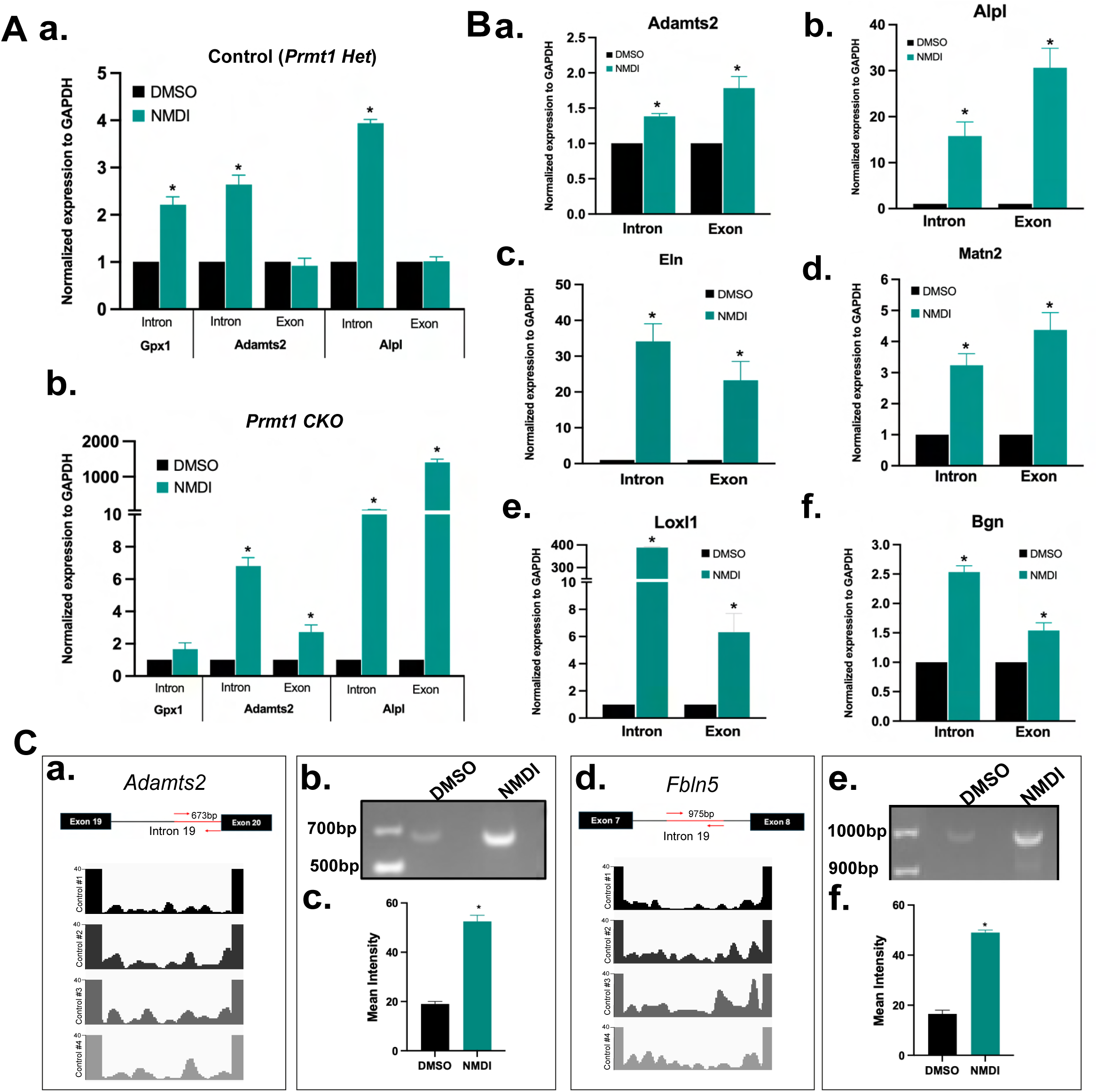
IR-triggered NMD functions as a basal and stress-responsive mechanism for mRNA decay in CNCCs. (Aa, Ab) Treatment with the NMD inhibitor NMDI14 (NMDI) led to accumulation of intron-retaining (intron) transcripts of *Gpx1*, *Adamts2*, and *Alpl* in CNCCs from Control (*Prmt1 Het)* and *Prmt1 CKO* embryos, analyzed by RT-PCR. Gpx1 serves as a positive control to validate NMD inhibition by NMDI14. (Ba–Bf) NMDI14 caused accumulation of intron-retaining (intron) and total mRNAs (exon) of *Adamts2*, *Alpl, Eln, Matn2, Loxl1* and *Bgn* in CNCCs. CNCCs were isolated from E13.5 *Wnt1-Cre; R26R^tdTomato^* and analyzed by RT-PCR. Primers that span the intronic or intron-exon junction region were used to assess intronic expression. Primers that span the exonic region were used to examine exonic expression that indicated mRNA abundance. (Ca, Cd) Intron retention of matrix transcripts *Adamts2* and *Fbln5* was detected in four independent control embryos by RNAseq, as illustrated by track views. (Cb-Cf) NMDI14 treatment caused accumulation of intron-retaining *Adamts2* and *Fbln5* transcripts. CNCCs isolated from E13.5 *Wnt1-Cre; R26R^tdTomato^* embryos were treated by DMSO or NMDI14 (NMDI), followed by mRNA extraction and assessment with semi-quantitative PCR. Primers (indicated by the red arrows) were designed to span regions (red line) of intron 19 of *Adamts2* or intron 7 of *Fbln5*. * p<0.05 NMDI vs. DMSO of the same group.

We noted significant accumulation of retained introns in control CNCCs from *Prmt1 Het* (Figure-4Aa), suggesting a possibility that IR-induced NMD is a basal or physiological mechanism to regulate mRNA levels in CNCCs. To assess this possibility, we isolated CNCCs from control *Wnt1-Cre; R26R ^tdTomato^* embryos at E13.5 and treated cells with NMDI14 or DMSO, followed by poly(A)+ mRNA extraction and RT-PCR analysis. Multiple ECM genes were assessed for the role of NMD, which revealed significant accumulation of retained introns within *Adamts2, Alpl, Eln, Matn2, Loxl1* and *Bgn*, accompanied by increased mRNA abundance following NMDI14 treatment (Figure-4Ba-Bf). To strengthen these findings, we used traditional semi-quantitative PCR to detect a longer span (600-1000bp) of the intronic region. In the NMDI14-treated group, accumulation of *Adamts2* intron 19 and *Fbln5* intron 7 were revealed by higher intensity bands compared to DMSO group (Figure-4Ca-4Cf). Overall, these findings demonstrate that the NMD process is responsible for the decay of ECM transcripts. More importantly, we defined IR-triggered NMD as a physiological mechanism in CNCCs to control ECM mRNA abundance and gene expression during craniofacial development. Upon molecular stress caused by *Prmt1* deletion, NMD is further exploited to disrupt ECM expression.

### PRMT1 methylates splicing factors SFPQ, EWSR1 and TAF15 in CNCCs

Next, we investigated the molecular mechanism by which *Prmt1* deletion disrupts intron retention. PRMT1 is a methyltransferase that catalyzes methylation of arginine (R) residues within RG/RGG/GAR repeats. These repeats are highly enriched in RNA-binding proteins, particularly splicing regulators (Thandapani et al., 2013). PRMT1 generates asymmetric dimethylation (ADMA) on arginine to regulate the turnover, subcellular localization, and activity of splicing regulators (Smith et al., 2004). We therefore hypothesized that PRMT1 regulates the retention of introns via methylation of splicing regulators. Earlier studies by Graham et al. and our team have characterized the landscape of arginine methylation governed by PRMT1 and identified a cohort of RNA processing proteins exhibiting PRMT1-dependent methylation (Hartel, Chew, Qin, Xu, & Graham, 2019). Among these, six splicing regulators are highly expressed in embryonic CNCCs: SFPQ, SRSF1, EWSR1, TAF15, TRA2B, and G3BP1 (Figure-5A). Given that pre-mRNA splicing occurs in the nucleus following transcription, we examined the subcellular localization of these splicing factors in control CNCCs from E13.5 mouse embryos. SFPQ, EWSR1 and TAF15 predominantly localized to the nucleus (Figure-5Ba-5Bl). TRA2B was distributed in both nucleus and cytoplasm (Figure-5Bm-p). While SRSF1 and G3BP1 were primarily expressed in the cytoplasm (Figure-5Bq-x). These observations were corroborated by quantitative analysis of the nuclear to cytoplasmic ratio of their expression (Figure-5C), suggesting nuclear proteins SFPQ, EWSR1, TAF15 and TRA2B as prime candidates to mediate PRMT1-controlled splicing activity. All four splicing factors are documented substrates for PRMT1-catalyzed methylation in Hela, Jeg3 or HEK293 epithelial cells (Hartel et al., 2019; Jobert, Argentini, & Tora, 2009; K. K. C. Li, Chau, & Lee, 2018; Pahlich, Bschir, Chiavi, Belyanskaya, & Gehring, 2005; Snijders et al., 2015). To determine whether PRMT1 is responsible for methylation of these splicing factors in the craniofacial structures, we employed proximity ligation assay (PLA), a technique designed for in-situ detection of protein modifications. To this end, E13.5 embryo sections were probed with anti-SFPQ antibody and anti-ADMA antibody that detects methyl-arginine using the PLA kit, which revealed the presence of methylated SFPQ as distinct green punctate staining. Within the craniofacial complex, SFPQ methylation was observed in the mandibular processes at E13.5 (Figure-5Da-5Dc, 5E). In *Prmt1* CKO embryos, the signal of methyl-SFPQ markedly decreased within the CNCC-derived mesenchyme region (Figure-5Dd-5Df, 5E). The same analysis was conducted in the maxillary regions, which also demonstrated robust SFPQ methylation in the control group and a dramatic reduction in *Prmt1*-deleted embryos (Figure-5Fa-5Ff, 5G). This CNCC-specific deletion of PRMT1 only reduced SFPQ methylation in the CNCCs, as non-CNCC lineages within the epithelium continued to display methyl-SFPQ signals (Supplemental Figure-S3A). To detect EWSR1 methylation, E13.5 embryo sections were probed with anti-EWSR1 antibody and anti-ADMA antibody using PLA kit. A low level of EWSR1 methylation was observed in both mandibular and maxilla processes at E13.5 (Figure-5H, 5I). Using the same approach, we also detected TRA2B methylation in mandibular and maxilla processes at E13.5 (Figure-5J, 5K), but the levels of EWSR1 and TRA2B methylation were much lower compared to SFPQ. In contrast to the low levels of methylation in craniofacial structures, the embryonic abdominal tissue showed prominent methylation of EWSR1 and TRA2B, validating the PLA techniques and EWSR1/TRA2B antibodies in efficient detection of their methylation (Supplemental Figure-S3B, S3C). We further determined whether methylation of EWSR1 and TRA2B depends on PRMT1 by comparing methylation signal between control and *Prmt1* deficient embryos and detected a significant reduction of methyl-EWSR1 and methyl-TRA2B within the CNCC-derived mesenchyme region in *Prmt1* CKO samples (Figure-5I-5K). TAF15 methylation wasn’t assessed due to lack of a compatible antibody. Collectively, these data indicate that SFPQ, EWSR1 and TRA2B are methylated by PRMT1 in the CNCCs at E13.5 within the craniofacial complex, with SFPQ showing the most prominent methylation and in a PRMT1-depedent manner.

**Figure 5.**
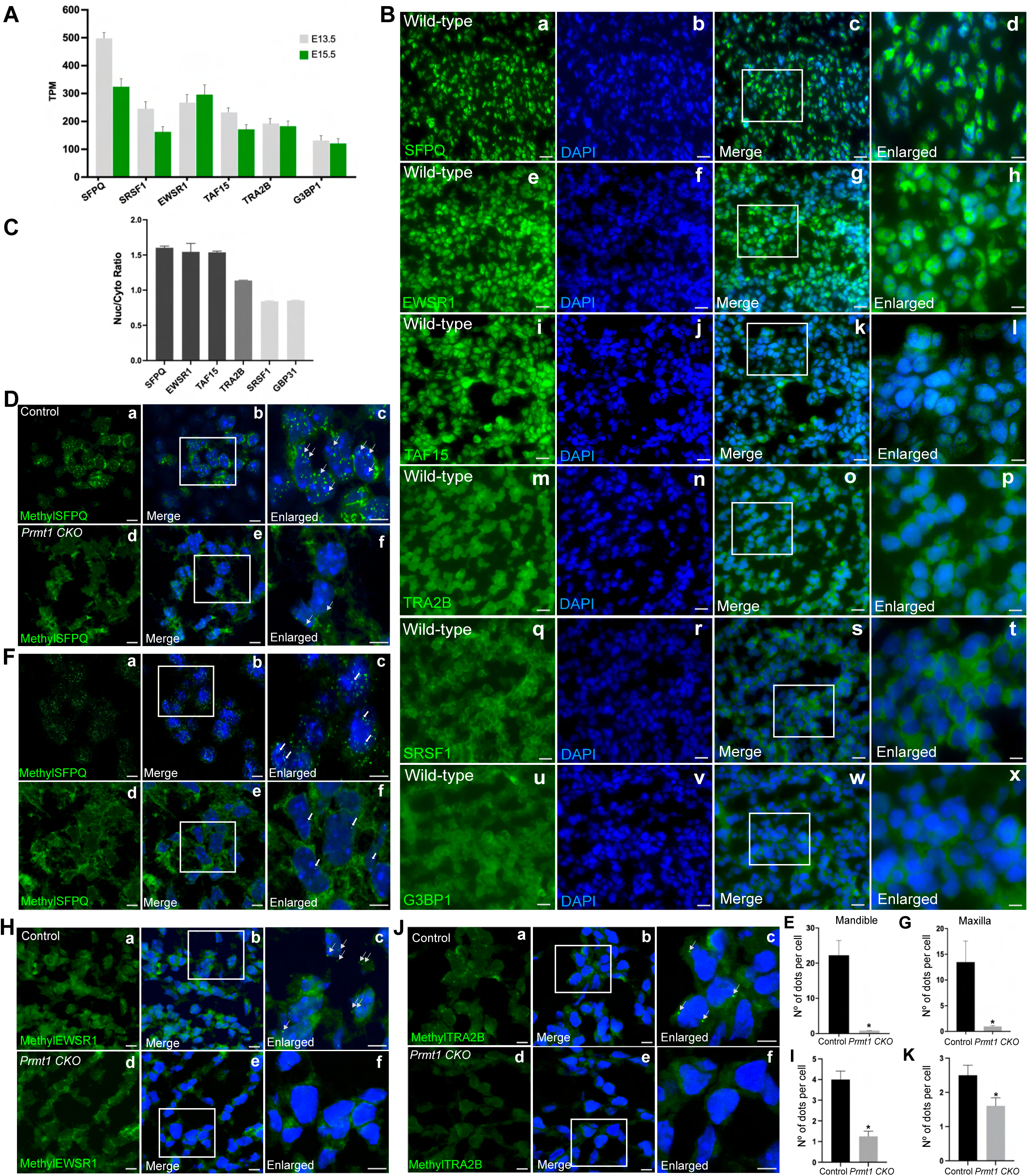
PRMT1 methylates SFPQ, EWSR1 and TRA2B in CNCCs. (A) Expression levels of splicing factors SFPQ, SRSF1, EWSR1, TAF15, TRA2B, HnRNPA1, WDR70 and G3BP1 in primary isolated CNCCs from *Wnt1-Cre; R26R^tdTomato^* mouse embryonic heads at E13.5 and E15.5. TPM, transcript per million. (B-C) Subcellular localization of splicing factors in CNCCs by immunostaining using sagittal sections of the mandibular process from wild-type mouse embryos, which revealed nuclear expression of SFPQ (a-d), EWSR1 (e-h), and TAF15 (i-l); even distribution of TRA2B between nucleus and cytosolic compartment (m-p), and cytoplasmic expression of SRSF1 (q-t) and G3BP1 (u-x). The subcellular distribution was quantified in C and presented as nuclear to cytosolic signal ratio (Nuc/Cyto Ratio). (D-G) SFPQ methylation diminished in the mandibular (D&E) and maxillary (F&G) processes of *Prmt1* deficient embryos at E13.5. (H-K) Reduction of EWSR1 and TRA2B methylation was observed in the mandibular processes of *Prmt1* deficient embryos at E13.5. Methylation was detected by proximity ligation assay (PLA). Green puncta indicated methyl-SFPQ, TRA2B or EWSR1. Nuclei were counterstained with DAPI (blue). Representative images are shown for Control (Da-Dc; Fa-Fc; Ha-Hc; Ja-Jc) and *Prmt1 CKO* (Dd-Df; Fd-Ff; Hd-Hf; Jd-Jf). Higher magnification views in (Dc, Df, Fc, Ff, Hc, Hf, Jc, Jf) illustrate methyl-SFPQ, TRA2B and EWSR1 (green puncta) in the nuclei, as indicated by white arrows. E, G, I and K showed quantification of PLA puncta normalized to cell number in four biological replicates, presented as mean ± SEM. * p<0.05 *Prmt1* CKO vs Control. Control: *Wnt1-Cre; R26R^tdTomato^*. *Prmt1 CKO: Wnt1-Cre; Prmt1^fl/fl^; R26R^tdTomato^*. Scale bar = 100µm in B, D, F, H and J except in enlarged panels, where scale bar = 25µm (Bd, Bh, Bl, Bp, Bt, Bx, Dc, Df, Fc, Ff, Hc, Hf, Jc, Jf).

### PRMT1-catalyzed methylation protects SFPQ from proteasomal degradation

PRMT1-induced methylation controls the activity of splicing factor by modifying their stability, subcellular distribution, or RNA binding affinity (Rho, Choi, Jung, & Im, 2007). We first focused on SFPQ and examined whether SFPQ protein expression or subcellular localization was disturbed by *Prmt1* deletion. In control embryos, SFPQ exhibited prominent expression in CNCCs and mainly localized in the nucleus within the mandibular and maxillary processes (Figure-6Aa-c, 6Ca-c, 5Ba-d, 5C). In *Prmt1*-deficient CNCCs, SFPQ protein expression level declined significantly in the maxilla and mandible (Figure-6A-D). A corroborating Western blot (WB) analysis using tissues from the craniofacial structures, including facial and anterior skull regions of embryonic heads, confirmed this decline in SFPQ expression (Figure-6E, 6F). These data suggest two possible mechanisms: PRMT1 promotes SFPQ expression or protects SFPQ from degradation. We first analyzed SFPQ mRNA levels in control and *Prmt1*-deficient CNCCs, which showed no discernable difference (Figure-6G), suggesting that loss of PRMT1 didn’t reduce SFPQ mRNA expression. Next, we tested whether PRMT1 regulates SFPQ degradation and assessed the proteosome-mediated degradation by treating CNCCs with MG132, a proteasome inhibitor. MG132 treatment in control CNCCs from *Prmt1 Het* embryos did not alter the expression of SFPQ, but in the *Prmt1* CKO CNCCs, MG132 caused a significant restoration of SFPQ protein (Figure-6H, 6I), suggesting that SFPQ protein in *Prmt1*-deficient CNCCs was reduced by proteasomal degradation. To test the relationship between arginine methylation and SFPQ degradation, we assessed SFPQ methylation in these MG132-treated CNCCs. SFPQ proteins rescued by MG132 in *Prmt1*-deficient CNCCs was not methylated (Figure-6J, 6K), suggesting that PRMT1-catalyzed arginine methylation may protect SFPQ from proteasomal degradation in CNCCs.

**Figure 6.**
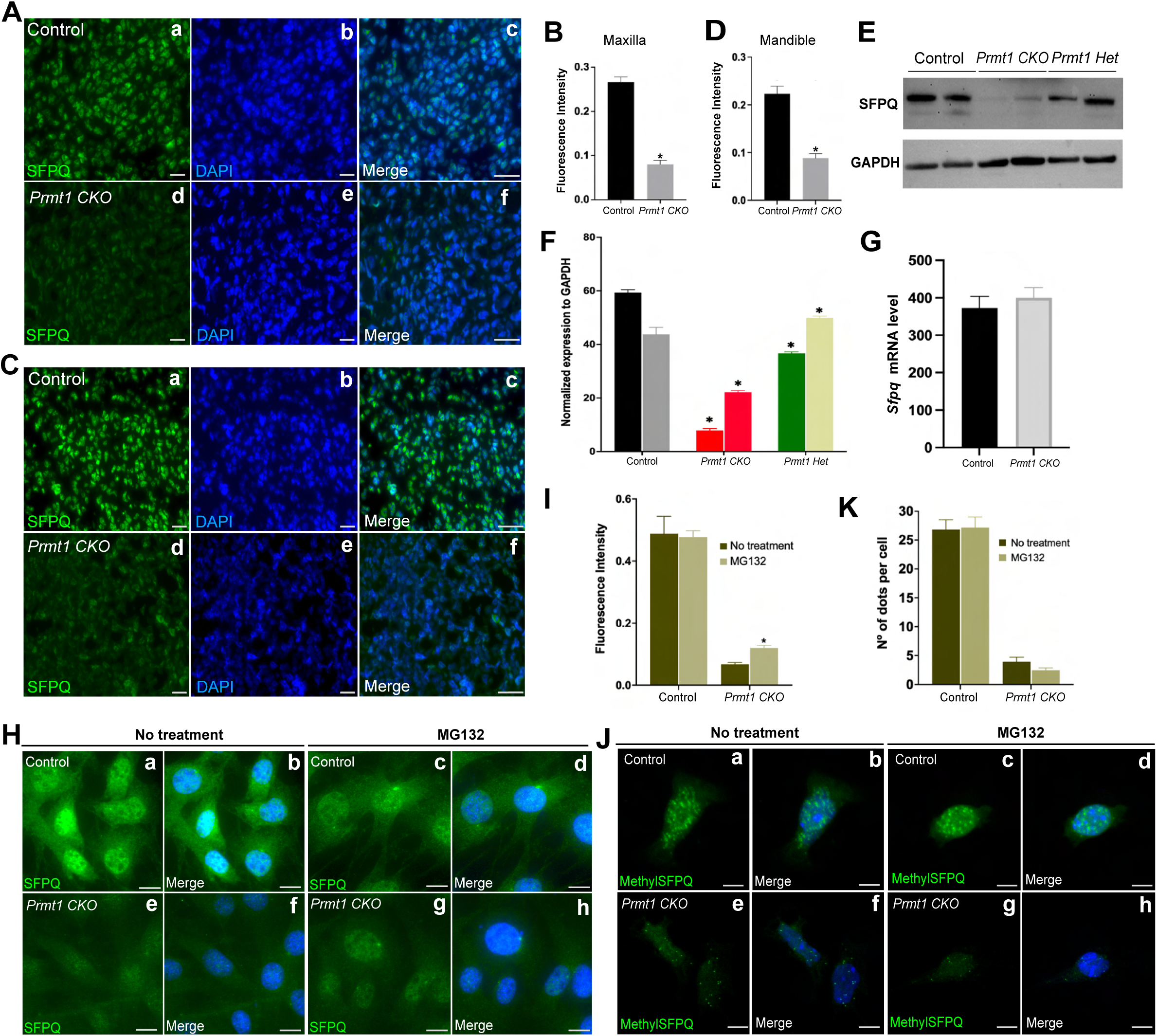
PRMT1 depletion reduces SFPQ protein levels via proteasomal degradation. (A-D) SFPQ protein level was significantly reduced in the mandibular (A&B) and maxillary (C&D) processes of *Prmt1* deficient embryos. SFPQ protein was detected by immunostaining and quantified in B and D. (E & F) SFPQ protein levels declined dramatically in the *Prmt1* deficient embryonic head, as detected by Western blotting (E) and quantified with ImageJ (F). (G) PRMT1 deletion did not alter SFPQ mRNA levels in CNCC. (H-K) SFPQ protein accumulated in *Prmt1-*deficient CNCCs upon MG132 treatment, and the accumulated SFPQ protein remained un-methylated. CNCCs isolated from *Prmt1 CKO* embryos and littermate controls (*Prmt1 heterozygous*) were treated with MG132. SFPQ protein was detected by immunostaining in H and quantified in I. SFPQ methylation was detected by PLA in J and quantified in K.* p<0.05 *Prmt1* CKO vs Control. Control, *Wnt1-Cre; R26R^tdTomato^*. *Prmt1 CKO, Wnt1-Cre; Prmt1^fl/fl^; R26R^tdTomato^*. Scale bar =100µm in A & C. Scale bar =25µm in H & J.

We further examine whether protein expression of EWSR1, TAF15 and TRA2B was disturbed by *Prmt1* deletion in the craniofacial complex. Unlike SFPQ, EWSR1, TAF15 and TRA2B showed a similar level of protein expression in *Prmt1* CKO and control embryos (Supplemental Figure-S4A-S4F). These findings were supported by WB analysis using tissues from the craniofacial structures (Supplemental Figure-S4G).

The subcellular distribution of EWSR1, TAF15 and TRA2B wasn’t altered either (Supplemental Figure-S4H-S4J). In summary, loss of *Prmt1* reduced SFPQ protein expression via proteasomal degradation, whereas the protein expression and subcellular localization of EWSR1, TAF15 and TRA2B were not altered by *Prmt1* deficiency.

### SFPQ regulates the splicing of matrix, Wnt signaling and neuronal genes in CNCCs

We further investigated the role of SFPQ in CNCCs. Primary CNCCs were isolated from control (*Wnt1-Cre; R26R^tdTomato^*) embryos at E13.5 and enriched by Td+ signal using cell sorting. Within a day, Td+ CNCCs were transfected with control siRNA or two independent SFPQ-targeting siRNAs to deplete SFPQ, and then poly(A)+ mRNA was purified followed by sequencing (Figure-7A). To determine whether these primary isolated CNCCs retained neural crest cell characteristics after siRNA-mediated knockdown, the expression of CNCC markers *Twist1, Sox10, Msx1, Snai2,* and *Tfap2a* was examined (Achilleos & Trainor, 2012; Ishii et al., 2012). The CNCC marker expression in transfected CNCCs after isolation is comparable to fresh CNCCs from E13.5 or E15.5 embryos (Supplemental Figure-S5), validating CNCC identity and supporting this method as a feasible approach for mechanistic study of neural crest cell biology using developmental stage-specific embryos. Subsequently, using sequencing data from these siRNA-transfected CNCCs which exhibit about 50% reduction of *Sfpq* (Figure-7A), we conducted bioinformatic analyses of differentially expressed genes (DEG) and differentially retained introns (IRI) between control and *Sfpq*-depleted CNCCs. First, we confirmed that the transcriptomic and intronic changes induced the two independent siRNAs targeting SFPQ are predominantly overlapping, as illustrated by scatter plot analysis of the DEG and IRI data (Figure-7Ba, 7Bb). Next, we focused on changes in retained introns caused by *Sfpq* depletion and revealed higher intron retention in 397 (siSFPQ #1 vs. siCont) and 388 (siSFPQ #2 vs. siCont) genes and lower IR in 129 (siSFPQ #1 vs. siCont) and 151 (siSFPQ #2 vs. siCont) genes (Figure-7Ca, Cb). Genes with higher IR also bear significant overlap with downregulated genes. Among 269 (siSFPQ #1 vs. siCont) and 232 (siSFPQ #2 vs. siCont) genes downregulated by *Sfpq* depletion, 81 (siSFPQ #1 vs. siCont) and 60 (siSFPQ #2 vs. siCont) genes overlapped with elevated IR, respectively (Figure-7Cc, Cd). *Sfpq* depletion also led to downregulation of 269 (siSFPQ #1 vs. siCont) and 232 (siSFPQ #2 vs. siCont) genes, among which 81 (siSFPQ #1 vs. siCont) and 60 (siSFPQ #2 vs. siCont) genes overlapped with elevated IR, respectively (Figure-7Cc, Cd). Genes showing higher intron retention in *Sfpq* depleted CNCCs encompassed “proteoglycan metabolic process” and “proteoglycan biosynthesis process”, which included GAG degradation genes, and “connective tissue development”, which includes ECM genes (Figure-7Da, 7Db). We also analyzed genes that were differentially expressed in control vs. *Sfpq*-depleted CNCCs (Figure-7E). GO analysis of differentially downregulated genes by SFPQ-specific siRNAs further revealed extracellular matrix and extracellular structure as the top pathways suppressed by SFPQ depletion, suggesting that IR-mediated regulation of ECM is a key functional consequence following *Sfpq* deficiency (Figure-7F). SFPQ has demonstrated direct roles in transcriptional regulation, where it facilitates RNA polymerase II (Pol II) recruitment and transcriptional termination in neuronal lineages (Takeuchi et al., 2018). To determine whether SFPQ directly regulates the transcription of matrix genes, we performed CUT&TAG with antibody again Rbp1, the largest subunit of Pol II, in ST2 cells, which is a mesenchymal cell line used for mechanistic studies of osteogenic differentiation (Canales et al., 2023; Ishida, Kawao, Mizukami, Takafuji, & Kaji, 2021; K. Li et al., 2019; Mizukami et al., 2023; Pregizer, Barski, Gersbach, García, & Frenkel, 2007; Seong et al., 2023; Strauss et al., 2019; Tu et al., 2007; Yang et al., 2019). ST2 cells were transfected with control or SFPQ-targeting siRNAs and collected for CUT&TAG followed by analyses of Pol II binding at the promoter region (3kb ± TSS) and gene body of matrix genes. Pol II recruitment was similar between control and SFPQ-depleted groups with marginal differences (Figure-7G, 7H), suggesting that transcriptional repression cannot fully account for the downregulation of matrix gene in SFPQ depleted groups.

**Figure 7.**
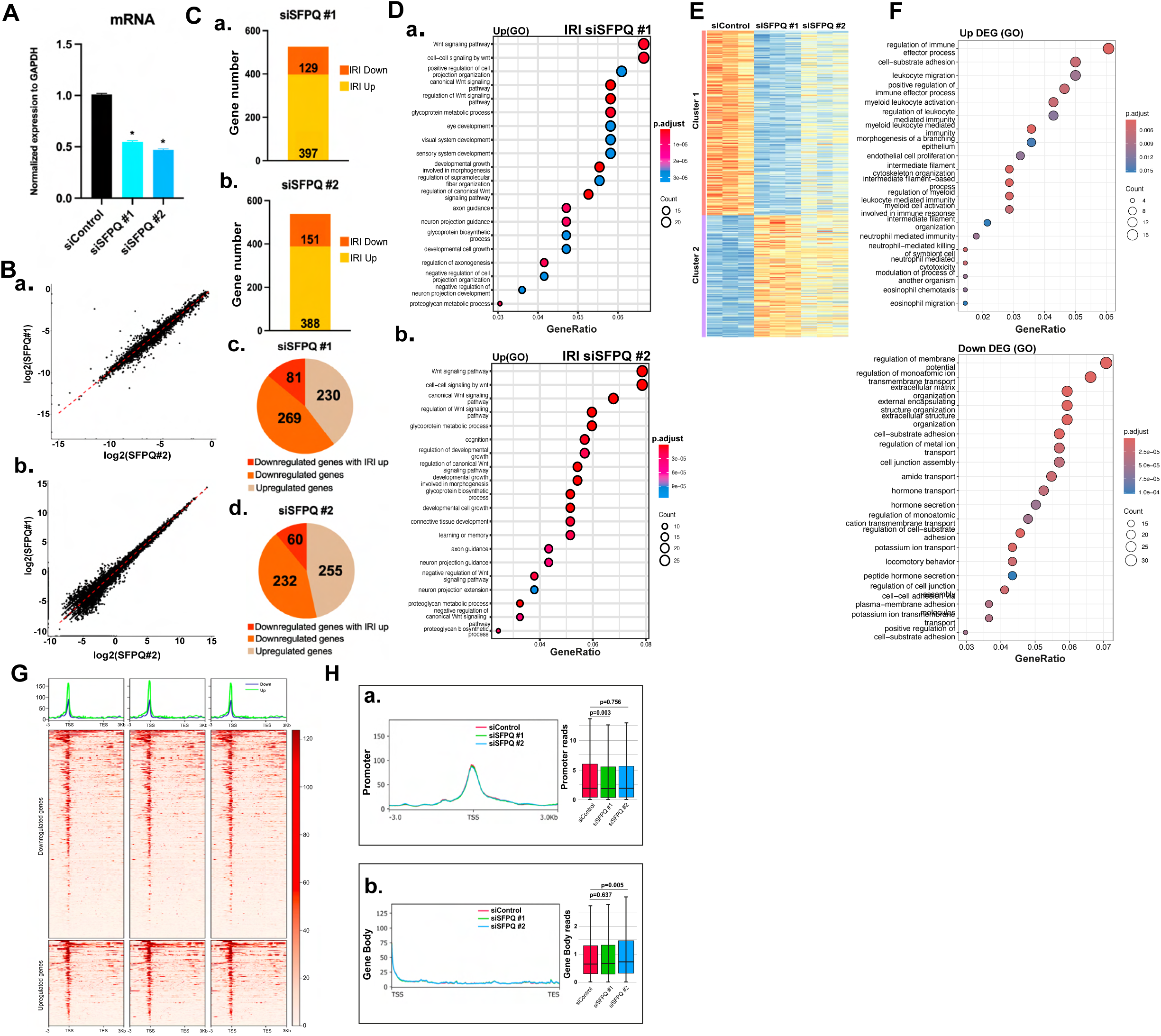
PRMT1-SFPQ pathway regulates matrix genes in CNCCs. (A) SFPQ knockdown caused around 50% reduction in *Sfpq* expression in CNCCs. CNCCs were transfected with two independent siRNAs targeting SFPQ, or control siRNA, followed by poly(A)+ mRNA extraction and RNA sequencing. * p<0.01 siSFPQ vs siControl. (B) The two independent siRNAs targeting SFPQ caused similar transcriptomic and intronic changes, shown by scatter plot of DEG (*Ba*) and IRI (*Bb*). DEG, differentially expressed genes. IRI, intron retention index. (C) SFPQ depletion altered intron retention in CNCCs. Changes of IR events in genes showing increased (Up, yellow color) or decreased (Down, orange color) IR were illustrated by stacked bar graphs. Pie chart demonstrated differentially regulated genes, with shaded areas among downregulated genes (orange) highlighting their overlap with IR elevated genes (red). (D) GO analysis of genes with elevated IR following SFPQ depletion. (E&F) Heatmap and GO analysis of SFPQ-regulated genes in CNCC. (G&H) Pol II CUT&Tag analysis in ST2 cells transfected with control or SFPQ siRNAs showing Pol II recruitment in downregulated and upregulated genes (*G*), promoter regions (*Ha*) and gene body (*Hb*). P-value was indicated at the top. siControl, control siRNA. siSFPQ#1 and siSFPQ#2, two independent SFPQ siRNAs.

More intriguingly, Wnt signaling emerged as the top pathway regulated by SFPQ in CNCCs (Figure-Figure-7Da, 7Db). *Sfpq* depletion increased the percentage of intron-retaining mRNAs of genes in this cluster and reduced total mRNA expression (Figure-8A). The Wnt signaling components-regulated by SFPQ encode both positive and negative regulators of Wnt signaling and encompass both canonical and non-canonical Wnt signal regulators. Additionally, neuronal genes constitute a big fraction among SFPQ-regulated introns as “axon guidance”, “axonogenesis”, “neuron projection development”, and “leaning or memory” emerged as the top biological processes among differentially upregulated introns and SFPQ-regulated genes, and “regulation of membrane potential”, “learning or memory” “cognition” and “locomotory behavior” denotes differentially downregulated genes (Figure-7F). In CNCCs, *Sfpq* depletion increased the percentage of intron-retaining mRNAs of these neuronal genes and reduced total mRNA expression (Figure-8B). Altogether, these findings demonstrate that depletion of SFPQ enhanced intron retention and reduced mRNA expression in matrix, Wnt signaling components and neuronal genes.

**Figure 8.**
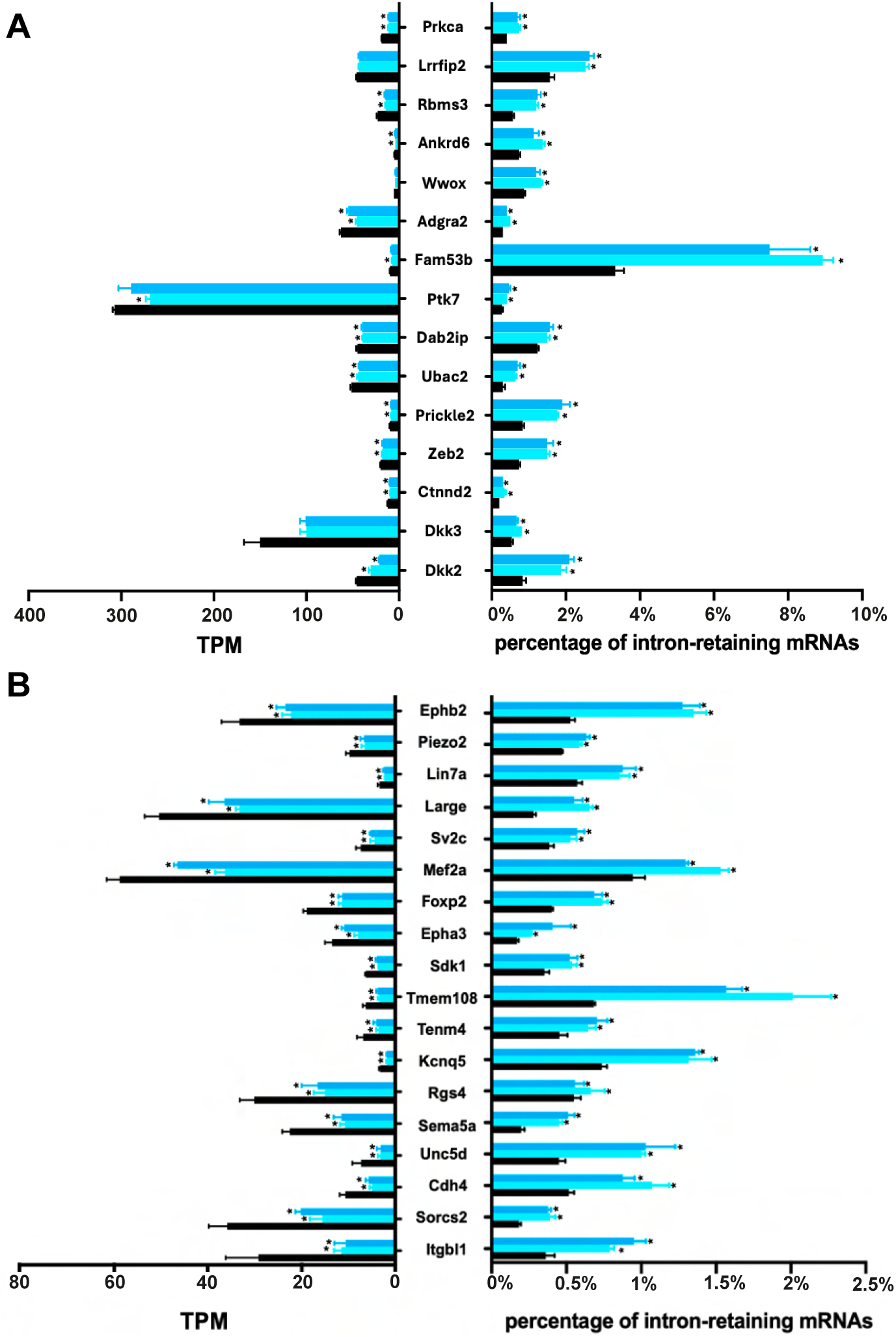
SFPQ regulates intron retention of Wnt signaling and neuronal genes in CNCCs. SFPQ depletion in CNCCs elevated intron retention and decreased mRNA abundance of Wnt signaling components (A) and neuronal genes (B). The levels of mRNA abundance (left) and intron retention (right) were illustrated by a two-sided bar graph. TPM, transcripts per million. * p<0.01 siSFPQ vs siControl. siControl, control siRNA. siSFPQ#1 and siSFPQ#2, two independent SFPQ siRNAs.

### SFPQ depletion reduces matrix and Wnt signaling gene expression via IR-induced NMD

In *Sfpq*-depleted CNCCs, retained introns also contained PTCs within their open reading frames, suggesting the involvement of PTC-triggered NMD (Supplemental Table S3). To investigate the role of IR-induced NMD, we utilized ST2 cells. SFPQ knockdown in ST2 cells caused an increase in retained introns and decrease in mRNA expression for *Col4a2, St6galnac3,* and *Ptk7* (Figure-9A). To determine whether *Col4a2, St6galnac3,* and *Ptk7* mRNA were degraded through IR-induced NMD, we inhibited NMD with NMDI14 (Martin et al., 2014). NMDI14 treatment caused accumulation of intron 4 of *Col4a2,* intron 1 of *St6galnac3,* and intron 1 of *Ptk7* (Figure-9B), indicating that the decay of intron-retaining transcripts in the *Sfpq*-depleted cells depends on the NMD machinery. To assess the impact of NMD on the abundance of these mRNAs, we designed PCR primers to detect the exonic regions and observed that inhibition of NMD restored and increased the abundance of these mRNAs in *Sfpq*-depleted ST2 cells (Figure-9C). These data indicate that depletion of *Sfpq* causes aberrant intron retention that triggers mRNA decay through NMD. We further noted that NMDI14 induced accumulation of intron 4 of *Col4a2* and intron 1 of *St6galnac3* in the control group (Figure-9B), accompanied by accumulation of their transcripts (Figure-9C), suggesting that NMD is also a basal mechanism to regulate gene expression in ST2 cells, a function similar as in embryonic CNCCs.

**Figure 9.**
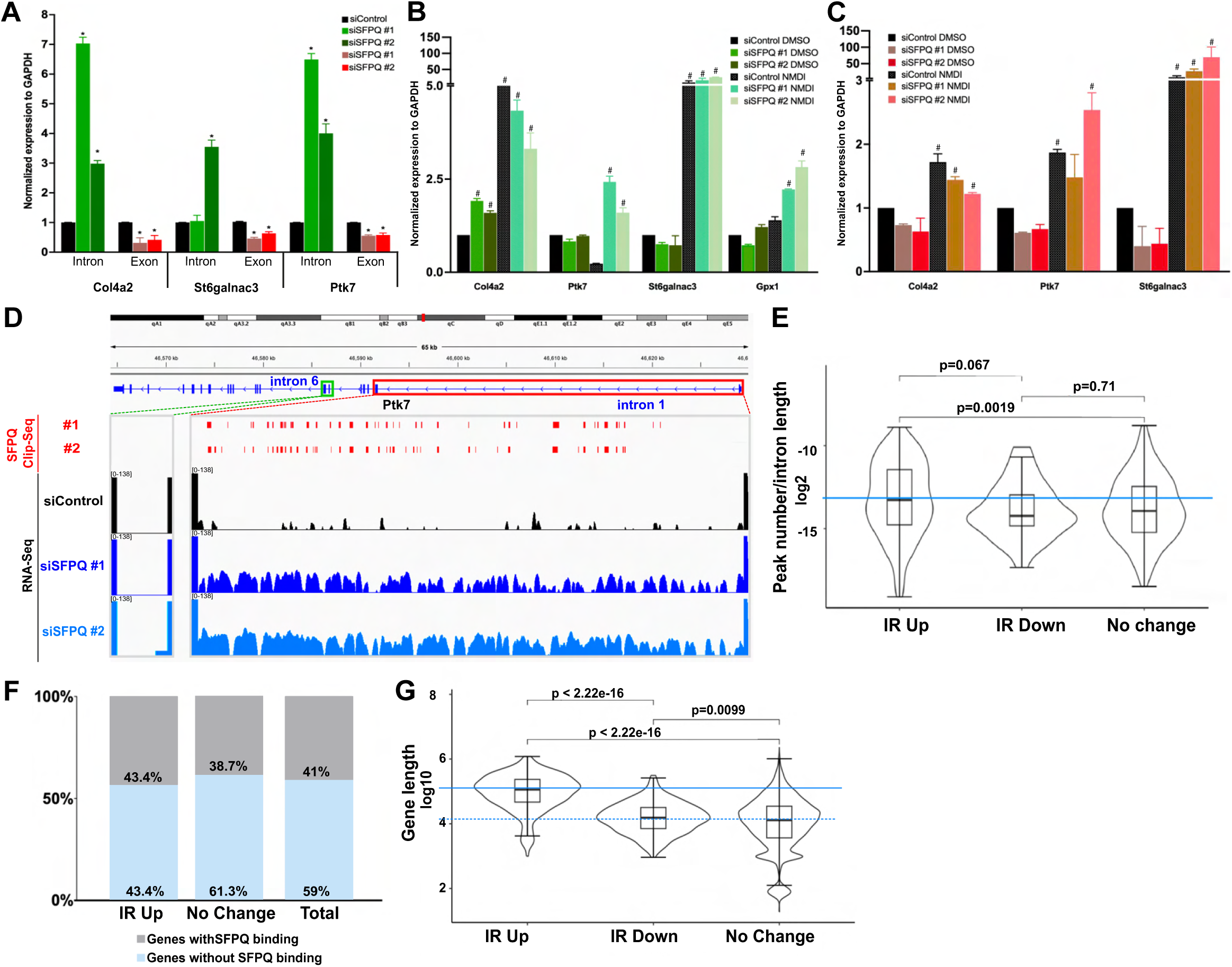
SFPQ depletion reduces long gene expression via intron retention triggered NMD. (A) SFPQ depletion promoted intron retention and reduced mRNA abundance of *Col4a2*, *St6galnac3*, and *Ptk7* in ST2 cells. Bar chart showing RT-PCR analysis of intronic and exonic expression. (B&C) NMD inhibitor NMDI14 caused the accumulation of retained introns and total mRNAs of *Col4a2, St6galnac3,* and *Ptk7* in ST2 cells. Bar chart showing RT-PCR analysis of intronic (B) and exonic expression (C) in DMSO or NMDI-treated cells. NMDL14-mediated inhibition of NMD was validated using Gpx1 as a positive control. *p<0.05 siSFPQ vs. siControl. ^#^ p<0.05 NMDI vs. DMSO treatment of the same group. (D) SFPQ binding peaks were mapped to retained intron 1 but not spliced intron 6 of *Ptk7*. Peak distribution from published *Sfpq* CLIP-seq data using E13.5 brain (top) and track view of RNA-seq data using siControl or siSFPQ-transfected CNCCs (bottom) for *Ptk7,* with the retained *Ptk7* intron 1 (red box) and spliced intron 6 (green box) highlighted. (E) SFPQ binding peaks were significantly enriched in retained intron regions within CNCCs. Violin plot displaying the density of SFPQ binding peaks in introns with elevated retention compared to introns with reduced retention or no change. P-value was calculated using Mann-Whitney U test and indicated at the top. (F) SFPQ binding peaks were preferentially enriched in genes with higher intron retention when compared to genes with no IR change. P = 0.07 using Fisher’s exact test. (G) SFPQ-regulated genes were significantly longer than average. Violin plot displaying the distribution of length for genes showing increased intron retention (IR Up), decreased intron retention (IR Down), or unchanged intron retention (No Change). Median length of the IR Up group was highlighted with solid blue line. Median length of the IR Down and No Change groups was highlighted with dotted blue line. P-value was calculated using Mann-Whitney U test and indicated at the top.

### Disturbance of SFPQ activity impaired splicing of long genes and long introns

SFPQ predominantly binds to intronic regions and regulates splicing (Hosokawa et al., 2019). We used published CLIP-seq dataset from embryonic mouse brain to assess whether these aberrantly retained introns are SFPQ targets (Hosokawa et al., 2019). As illustrated in *Ptk7* sequence, the retained intron 1 exhibited a much higher density of SFPQ binding peaks when compared to the constitutively spliced intron 6 within the same gene (Figure-9D). We further conducted analysis for all introns at the genomic level by comparing SFPQ binding peaks among introns with higher retention to introns with unaltered or lower retention. The comparison demonstrated that introns with elevated retention in *Sfpq*-depleted CNCCs showed significantly higher enrichment of SFPQ binding peaks (Figure-9E). We also analyzed SFPQ CLIP-seq peaks at the gene level, comparing genes with elevated IR to genes with no IR changes and showed that genes containing elevated IR in *Sfpq*-depleted CNCCs exhibited a trend towards higher enrichment of SFPQ binding peaks based on this embryonic brain dataset (Figure-9F). These data suggest that introns with aberrant IR elevation in *Sfpq*-depleted CNCCs are splicing targets bound by SFPQ.

To understand how SFPQ targets these matrix, Wnt and neuronal genes and introns, we examined molecular characteristics shared among SFPQ-regulated genes and recognized that they are long genes (>100kd in length) and retain long introns (>10kd in length). Of note, Wnt pathway gene *Wwox* and matrix genes *St6galnac3* are 913kd and 526kd in length, and their retained introns are 639kd and 215kd in length, respectively, in contrast to a median size of 0.6 ∼ 2.4kb for introns in the mouse genome (Hong, Scofield, & Lynch, 2006). In the human and mouse genome, long genes mostly occur because of long introns (Breschi, Gingeras, & Guigó, 2017; Lopes, Altab, Raina, & de Magalhães, 2021). We then plotted the length of SFPQ-regulated genes against genes across the genome and demonstrated that the genes with increased IR in *Sfpq*-depleted CNCCs have a median length of ∼100kb. This is much longer than gene length in the control groups, where the medial length is around 10kb (Figure-9G). These findings suggest that SFPQ-regulated splicing facilitates regulation of long gene expression during craniofacial development.

### SFPQ, EWSR1, TAF15 and TRA2B regulates distinct transcriptional and splicing events

To assess genes and IR co-regulated by SFPQ and PRMT1, we cross-analyzed genes with intron retention in both *Sfpq*-depleted CNCCs and *Prmt1*-deficient CNCCs. In *Sfpq*-depleted CNCCs, 179 genes exhibited significantly higher IR in both siRNA groups. In *Prmt1*-deficient CNCCs, 773 genes exhibited significantly higher IR in all embryos analyzed (Figure-10A). The cross-analysis showed that SFPQ-regulated intron retention bears partial (64 out of 773) overlap with PRMT1-regulated introns, exemplified by matrix genes *Col4a2, Adam12, Ntn1, App, St6galnac3, Galnt11 Galnt10,* and *Asph* (Figure-10A-E). Wnt signaling genes *Wwox* and *Dkks*, and neuronal genes that regulate membrane potential (38 out of 303 genes) was also identified among genes with decreased expression in *Prmt1 CKO* CNCCs, suggesting the regulation of these genes by the PRMT1-SFPQ pathway. The overlap may be under-represented because SFPQ was reduced to 50% by the depletion, but the partial overlay does suggest a notion that PRMT1 regulates intron retention through multiple splicing factors.

**Figure 10.**
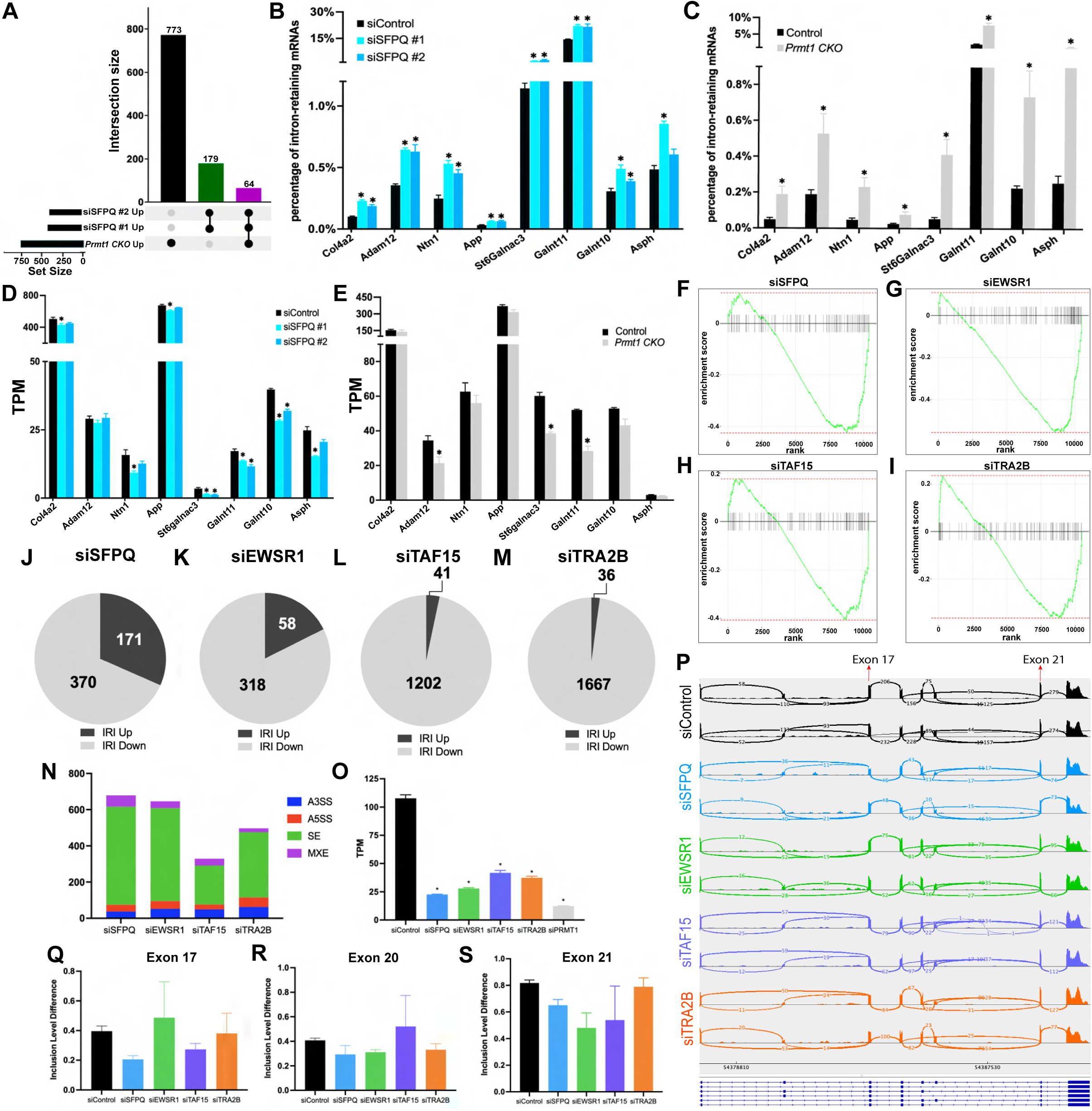
SFPQ, EWSR1, TAF15, and TRA2B regulate distinct transcriptional and splicing programs. (A) Genes with increased IR events in SFPQ-depleted CNCCs demonstrated 8.28% (64 out of 773) overlap with *Prmt1* deficient CNCCs. (B-E) Matrix genes *Col4a2, Adam12, Ntn1, App, St6Galnac3, Galnt10* and *Asph* were regulated by both *Prmt1* deletion and SFPQ depletion. Bar graph showing elevated IR (*B, C*) and reduced mRNA abundance (*D, E*) in CNCCs. TPM, transcripts per million. * p<0.05 siSFPQ vs siControl. * p<0.05 *Prmt1 CKO* vs Control. Control: *Wnt1-Cre; R26R^tdTomato^*. *Prmt1 CKO: Wnt1-Cre; Prmt1^fl/fl^; R26R^tdTomato^*. (F-S) ST2 cells were transfected with control or SFPQ, EWSR1, TAF15, TRA2B siRNAs and mature mRNA was extracted for sequencing. GSEA demonstrated enrichment of ECM genes among downregulated genes upon depletion of SFPQ, EWSR1, TRA2B, and TAF15 (*F-I*). Pie charts showed the percentage of genes with increased (dark grey) or decreased IRI (light grey) upon SFPQ, EWSR1, TAF15 and TRA2B deletion (*J-M*). rMATS analysis showed widespread and significant splicing changes following depletion of SFPQ, EWSR1, TAF15, and TRA2B (*N*). *Postn* mRNA expression was decreased in all four depletion groups (*O*). Exon skipping events of *Postn* were noted in all four depletion groups, illustrated by Sashimi plots (*P*) and quantified by bar graphs based on rMATS analysis (*Q-S*). * p<0.05 siRNAs vs siControl. siControl, control siRNA. siSFPQ#1 and siSFPQ#2, two independent SFPQ siRNAs.

We further characterized gene expression and intron retention regulated by SFPQ, EWSR1, TAF15 and TRA2B using ST2 cells. To this end, cells were transfected with control siRNA or two independent siRNAs targeting each splicing factor, and then poly(A)+ mRNA was purified followed by sequencing. GSEA analysis demonstrated that depletion of SFPQ, EWSR1, TRA2B, and TAF15 each caused downregulation of ECM genes (Figure-10F-I), suggesting that the four splicing factors all promote ECM gene expression. Global analysis for differentially expressed genes further showed that these splicing regulators control distinct biological functions (Supplemental Figure-S6). We also analyzed intron retention following their knockdown. siSFPQ caused differential IR in 514 genes, with increased IR in 171 genes and decreased IR in 370 genes (Figure-10J, Supplemental Table S4). siEWSR1 caused increased IR in 58 genes and decreased IR in 318 genes (Figure-10K, Supplemental Table S4), where genes with elevated IR belonged to diverse biological functions (Supplemental Table S5). In contrast, siTAF15 and siTRA2B caused predominantly decrease in IR with IR increase in <3% of genes with differential IR (Figure-10L-M, Supplemental Table S4). These findings suggest that the roles of EWSR1, TAF15 and TRA2B are distinct from SFPQ in the regulation of IR. Bioinformatic analysis for EWSR1, TAF15 and TRA2B didn’t reveal discernable overlap between ECM genes and IR changes, suggesting that they regulate ECM gene expression through IR-independent mechanisms in ST2 cells. Examination of other types of alternative splicing events by rMATS analysis (multivariant analysis of transcript splicing, MATS) revealed many splicing changes (Figure-10N, Supplemental Table S6). For example, the ECM gene *Postn* is known to exhibit exon skipping at *exon 17* and *21,* which affects *Postn* expression and activity that alter mandibular morphogenesis (Ishihara et al., 2023). We observed that the mRNA expression of *Postn* is reduced in all four splicing factor depletion groups and exon skipping of *Postn* showed a trend towards enhancement by the depletion, as illustrated by Sashimi plots and quantified from rMATS analysis (Figure-10O-10S).

To identify additional splicing regulators accountable for IR changes in *Prmt1* deficient embryos, we conducted in-silico analysis using RNA-seq data of mandibular CNCCs from control and *Prmt1 CKO* embryos by performing rMAPS2 (RNA map analysis) subsequent to rMATS (Hwang et al., 2020; Park, Jung, Rouchka, Tseng, & Xing, 2016; Y. Wang et al., 2024). A list of RNA-binding proteins (RBPs) was identified with highly significant binding scores in the differentially expressed genes, suggesting them as potential downstream mediators for PRMT1-regulated splicing changes. This in-silico approach predicted additional splicing regulators that may function downstream of PRMT1 in elevating intron retention, including SRSF9, Lin28A, PABPN1, ESRP2, TARDBP, KHDRBS1 and FXR1 (Supplemental Table S7). SFPQ is also among the RBPs with top occurrences, with SFPQ binding motifs depicted in the vicinity of alternative splicing events (Supplemental Figure-S7A-S7E, Supplemental Table S8). Taken together, PRMT1-regulated splicing in CNCCs is mediated by a plexus of splicing factors including SFPQ.

## DISCUSSION

In this study, we revealed prevalent intron retention within CNCCs during embryonic development and demonstrated that PRMT1-SFPQ pathway regulates the splicing of matrix and Wnt pathway genes during craniofacial development. Genetic deletion of *Prmt1* or depletion of *Sfpq* in CNCCs caused aberrant IR that triggers NMD to degrade intron-retaining transcripts. To our knowledge, this study is the first to characterize functional significance of IR-induced NMD in CNCCs and during craniofacial development. A main feature shared by SFPQ-regulated genes in CNCCs is gene length, with a median length of 100kb, in contrast to the median length of ∼10kb across the genome. Introns exhibiting high retention also tend to be long introns, for example, intron 4 of *Col4a2* which displayed dramatically increased retention in *Sfpq* deficiency boasts a length of 45kb. Long genes with long introns are linked to neuronal function, where retained introns have been proposed to facilitate a rapid response to external stimuli, allowing a time frame shorter than that required for de novo transcription (Mauger, Lemoine, & Scheiffele, 2016; Ni et al., 2016).

Genes over 100kb in length require two to several hours for transcription based on a rate of 1-2kb/minute, which poses a challenge for the fast tempo of embryogenesis. This intron-regulated mechanism may represent a physiological strategy that enables rapid production of matrix proteins when they are required for craniofacial morphogenesis. This post-transcriptional regulation allows cells to control the rate of transcript production, and to achieve rapid protein translation without waiting for the time period required for transcription. Long genes regulated by this mechanism are highly enriched in matrix formation, responsible for the production of building blocks for bone, cartilage, and connective tissues. Since spliceosomopathies preferentially affect the craniofacial skeleton and limb, IR-regulated matrix expression provides mechanistic insights for the higher susceptibility of these tissue types to spliceosome dysfunction. Many of the PRMT1-SFPQ pathway regulated matrix and Wnt signaling genes are associated with congenital defects. For example, in our study, the long gene *Wwox* exhibited increased retention of long introns (introns 3 and 4) and decreased expression upon *Prmt1* or *Sfpq* deficiency. In human patients, pathogenic variants of *WWOX* with large deletion within the long introns have been associated with epileptic encephalopathy syndrome manifesting shared facial phenotype (Dvinge & Bradley, 2015). The deletion within these long introns spanning the SFPQ binding region may affect SFPQ recognition and subsequent splicing of *WWOX* transcripts. Variants of another SFPQ target, *Ptk7,* are associated with neural tube defects and scoliosis (Berger, Wodarz, & Borchers, 2017). *SFPQ* itself is also linked to neurodegenerative diseases including amyotrophic lateral sclerosis, frontotemporal dementia, and Alzheimer’s disease, and was recently identified as a genetic regulator in CNCCs through an integrated genomic analysis (Feng et al., 2021). Besides SFPQ-regulated targets, splicing factors we identified in the unbiased in-silico analysis, such as Rbfox2 and HuR (Supplemental Table S5), are also associated with craniofacial defects in human patients and mouse studies (Cibi et al., 2019; Glessner et al., 2014; Homsy et al., 2015; McKean et al., 2016; Verma et al., 2022). We propose that IR-induced downregulation acts as a causal mechanism for dysregulated gene expression that contributes to craniofacial deformity.

IR-triggered NMD in controlling the expression of matrix genes in CNCCs presented a distinct mechanism in CNCCs. NMD is the RNA turnover pathway to degrade aberrant RNAs, driven by a protein complex composed of SMG and UPF proteins(Han et al., 2018; Tan, Stupack, & Wilkinson, 2022). Intron retention-triggered NMD in cancer has both promotive and suppressive roles and NMD inhibitors has been tested for cancer therapy and recently cancer immunotherapy(Tan et al., 2022). During embryonic development, the functional significance of NMD machinery is implied by human genetic findings and mouse genetic models. *SMG9* mutation in human patients causes malformation in the face, hand, heart and brain (Shaheen et al., 2016). *Smg6, Upf1, Upf2* and *Upf3a* knockout mouse die at early embryonic stages (E5.5-E9.5), and *Smg1* gene trap mutant mice die at E12.5, all presenting severe developmental defects (Han et al., 2018). In this study, upon *Prmt1* or *Sfpq* deficiency, IR-induced NMD adopted a pathogenic role, degrading aberrant intron-retaining transcript to reduce gene expression. We also demonstrated a physiological role for IR-triggered NMD that tunes matrix gene expression during embryogenesis, as inhibition of NMD using chemical inhibitors led to accumulation of transcripts such as *Alpl, Eln and Loxl1*.

In mammalian cells, PRMT1 methylates over forty splicing factors to modulate protein stability, subcellular localization, thereby influencing their accessibility to mRNA (Thandapani et al., 2013; Wu et al., 2022). PRMT1-catalyzed arginine methylation also affects their RNA binding affinity and specificity, altering their activity and the splicing product (Fedoriw et al., 2019; Fong et al., 2019). Therefore, it is not surprising that the disruption of this methyltransferase leads to splicing defects. We recognized SFPQ as a key PRMT1 substrate in CNCCs and demonstrated abundant SFPQ methylation in the mandibular and maxillary primordium. Our previous work also revealed SFPQ methylation on arginine (Arg) 7 and 9 depends on PRMT1(Hartel et al., 2019). This aligns with previous work demonstrating that PRMT1 is responsible for the association of SFPQ with mRNA in mRNP complexes in mammalian cells (Snijders et al., 2015). Our findings on altered neuronal genes in *Sfpq-*depleted CNCCs also echoed previous reports in which SFPQ regulates many neuronal genes involved in axon extension, branching, viability, and synaptogenesis via direct binding to their intronic regions (Cosker, Fenstermacher, Pazyra-Murphy, Elliott, & Segal, 2016; Ruskin, Zamore, & Green, 1988). Loss of SFPQ function compromises splicing patterns, especially the accurate splicing of long introns in embryonic brain and AML samples (Luisier et al., 2018; Taylor et al., 2022; Thomas-Jinu et al., 2017). Findings in our study further revealed previously unappreciated roles of SFPQ in the regulation of matrix genes through IR, which bear functional significance in bone and cartilage matrix deposition and craniofacial skeleton formation. Manipulation of SFPQ expression (knockdown with 50% efficiency) demonstrated that SFPQ shares around a partial overlap with PRMT1 in IR-regulated neural crest gene expression. Given that PRMT1 methylates over 40 splicing factors in mammals and many of these splicing factors are abundantly expressed in embryonic CNCCs, we expect that PRMT1-regulated intron retention will be mediated by multiple splicing regulators besides SFPQ. This notion is further supported by the identification of additional splicing regulators via in-silico analysis. Additionally, PRMT1-governed splicing will be mediated by multiple RNA binding protein encompassing all types of alternative splicing, as suggested by PRMT1-mediated methylation of EWSR1 and TRA2B, and the distinct splicing footprints among EWSR1, TAF15, TRA2B and SFPQ.

Altogether, these findings demonstrate that the PRMT1-SFPQ pathway regulates IR in CNCCs and suggests IR-triggered NMD as a mechanism that controls matrix and Wnt signaling gene expression during craniofacial morphogenesis.

## MATERIAL AND METHODS

### Animals

*Prmt1fl/fl* mice were generously provided by Dr. Stephane Richard (McGill University). *Wnt1-Cre; Prmt1^fl/fl^; R26R ^tdTomato^* and *Wnt1-Cre; R26R^tdTomato^* mice were obtained by mating *Wnt1-Cre, R26R^tdTomato^* mice (Jax #009107 and #007914) with *Prmt1fl/fl* mice. Animals were genotyped using established protocols (Chai et al., 2000; Yu et al., 2009) and all care and experiments followed USC’s IACUC protocols.

### Proximity Ligation Assay (PLA)

PLA was performed using a Duolink In Situ kit (Millipore Sigma Cat# DUO92101-1KT), SFPQ antibody (Everest Biotech Cat#EB09523), and pan-methyl arginine antibody, Asymmetric Di-Methyl Arginine (ADMA) (Cell Signaling Technology Cat# 13522) to detect methyl-SFPQ. Tissue sections were blocked with the Duolink blocking solution in a humidity chamber for 1 hour at 37°C before incubating with the primary antibodies (diluted 1:50 in Duolink PLA probe diluent) overnight at 4°C. The next day, tissue sections were incubated with the Goat-PLUS and Rabbit-MINUS PLA probes for 1 hour at 37°C, followed by ligase and amplification solution. After the final washes, tissue sections were mounted with Duolink In Situ mounting media with DAPI to counterstain nuclei. Images were visualized under the Confocal microscope. The number of PLA signals that appear as punctuate green signals in the nuclei of the cells and the quantification was performed with CellProfille software.

### Tissue Processing and Immunofluorescence Staining

E13.5 control and *Wnt1-Cre; Prmt1^fl/fl^* mice embryos were fixed in 4% PFA overnight, dehydrated, embedded in O.C.T., sectioned at 8 µm thickness in sagittal orientation, and mounted on glass slides. The samples were submitted to antigen retrieval using Antigen Unmasking Solutions (Vector Laboratories Cat# H-3300-250) and washed and incubated in 0.5% Triton X-100 in PBS for 20 minutes for permeabilization. The samples were briefly washed with PBS, blocked with 10% goat serum for 1 hour, and then incubated with primary antibody SFPQ (Cell Signaling Technology Cat#23020, 1:100) at 4°C overnight in a moisture chamber. After this, the samples were blocked with a secondary antibody (1:400, goat anti-rabbit IgG (H+L) Alexa Fluor 488 Cat# A-11008) for 1 hour followed by DAPI diluted 1:1000 in PBS for 10 minutes. Finally, the samples were mounted with mounting media (Electron Microscopy Science Cat# 1798510) and covered with cover glass for imaging using Keyence BZ-X800 and Leica DMI 3000B microscopes and quantification with CellProfille software.

### Western Blotting

Heads of E13.5 control and *Wnt1-Cre; Prmt1^fl/fl^* mice were dissected, followed by removal of the brain. The tissue was homogenized with a pestle followed by lysis with RIPA Lysis and Extraction Buffer. The total protein concentration was determined by comparison with BSA standards. Twenty micrograms of total protein from each sample were loaded into each well of a 10% polyacrylamide gel. Western blot analysis was carried out as previously described (Iwata et al., 2012). Antibodies against PRMT1 (Cell Signaling, Cat# 2449; 1:1000), SFPQ (Cell Signaling Technology Cat#23020, 1:1000), and GAPDH (Cell Signaling, Cat# 97166; 1:5000) were used for Western blot. The samples were analyzed three times in independent blots and the bands were quantified by optical densitometry. The analysis was performed using ImageJ digital imaging processing software (ImageJ 1.48v, National Institutes of Health, Bethesda, MD, USA). The expression of each analyzed protein was normalized with GAPDH.

### NCC collection and FACS analysis

The heads of E13.5 *Wnt1-Cre; R26R^tdTomato^* and *Wnt1-Cre; Prmt1^fl/fl^;R26R^tdTomato^* mice were dissected and placed in a sterile tube containing Hanks’ Balanced Salt Solution (Thermo Fisher Scientific Cat# 14175079). Following centrifugation, the supernatant was discarded, and the tissue was subjected to TrypLE (Thermo Fisher Scientific Cat# 12605010) incubation for 5-15 minutes at 37°C on a rotator. The resulting dissociated tissue was neutralized by the addition of fetal bovine serum (FBS) and then filtered through a 40μm strainer to obtain a single-cell suspension. After centrifugation, cells were resuspended in an appropriate volume of serum-free medium for FACS analysis. Sorting was performed based on tdTomato fluorescence (Excitation/emission: 554/582 nm), and td-positive cells were collected into separate tubes containing DMEM (Genesee Scientific Cat#25-501) supplemented with 20% FBS.

### Primary CNCC culture and siRNA transfection

CNCCs were cultured in DMEM supplemented with 20% FBS in an incubator until they attach to the bottom. Then, reverse transfection with control siRNA (Qiagen Cat# 1027310), SFPQ siRNA #1 (Qiagen Cat#SI05783848), and SFPQ siRNA #2 (Qiagen Cat#SI05783876) at 40 nM using Lipofectamine RNAiMax transfection reagent (Invitrogen) for siRNA delivery were performed. 48h later, total RNA was isolated using TRIzol reagent (Invitrogen) following the manufacturer’s protocols followed by mRNA isolation using NEBNext High-Input Poly(A) mRNA Magnetic Isolation Module (NEBNext® Cat#E3370S).

### mRNA isolation and sequencing

Poly(A) mRNA isolation was extracted from total RNA by using the NEBNext High-Input Poly(A) mRNA Magnetic Isolation Module (NEBNext® Cat#E3370S). Five sets of primary isolated CNCCs from control (*Wnt1-Cre;R26R ^tdTomato^)* or *Prmt1 CKO* (*Wnt1-Cre; Prmt1^fl/fl^; R26R ^tdTomato^),* and three sets of isolated CNCCs transfected with siControl or siSFPQs were sequenced at 40 million reads sequencing depth and 150 bp paired-end sequencing.

### Bioinformatic Analysis of Differentially Regulated Genes (DEG) and differential Intron Retention (IR)

The sequencing quality of RNA-seq libraries was assessed by FastQC v0.11.8 (https://www.bioinformatics.babraham.ac.uk/projects/fastqc/). Low-quality bases and adapters were trimmed using Trim Galore (v0.6.7,https://www.bioinformatics.babraham.ac.uk/projects/trim_galore/). The reads were mapped to mouse genome mm10 using hisat2 (v2.1.0) (Kim, Paggi, Park, Bennett, & Salzberg, 2019). The mapped sam files from hisat2 were converted to bam files which were then further turned to sorted bam files by samtools (v1.7) (H. Li et al., 2009). Mapped reads were then processed by htseq-count (v1.99.2) to calculate the read count of all genes (Anders, Pyl, & Huber, 2015). The sorted bam files were then given to IRTools (https://github.com/WeiqunPengLab/IRTools/) to calculate the IR of each gene and the read count of genes’ constitutive intronic regions (CIRs) and constitutive exonic regions (CERs). The expression level of a gene was expressed as TPM (Transcripts Per Kilobase Million) value. EdgeR was used to identify differentially expressed genes (DEGs) by requiring ≥ 1.5-fold expression changes and adjusted p-value < 0.05 (Robinson, McCarthy, & Smyth, 2010). The intron retention index (IRI) of a gene is defined as the ratio of the overall read density of its CIRs to the overall read density of its CERs. Genes with 0< IRI <1 and CER expression greater than 1 were selected to identify differential IR genes. Student’s t-test was used to identify differential IR genes by requiring ≥ 1.5-fold log2(IR) changes and p-value < 0.05. Gene Ontology analysis was done by DAVID (https://david.ncifcrf.gov/tools.jsp) and Metascape (https://metascape.org/gp/index.html#/main/step1) (Sherman et al., 2022; Zhou et al., 2019).

The RNA-seq data are deposited to the Gene Expression Omnibus (GEO) database with accession number GSE266474.

### rMATS and rMAPS Analysis

rMATS (https://github.com/Xinglab/rmats-turbo) and rMAPS2 (http://rmaps.cecsresearch.org/) were used to find potential RNA binding Proteins (RBPs) that contribute to differential splicing events (Hwang et al., 2020; Y. Wang et al., 2024). rMATS was first used to identify differential alternative splicing events between wild type and PRMT1 CKO samples by requiring FDR < 0.05. The output of rMATs were then fed to rMAPS2 to identify significant RBPs contributing to each type of splicing events.

### Enrichment analysis of SFPQ binding

To assess the SFPQ binding of the genes whose IR is regulated by SFPQ, we downloaded SFPQ CLIP-seq peak data from GEO (GSE96080). The CLIP-seq peaks shared between the replicates were filtered by p < 0.01 and log(FC) > 1 to obtain the significant peaks for downstream analysis. Next, we defined the SFPQ-regulated IR up and down gene sets by combining those from two SFPQ knockdown conditions respectively. We constructed the control gene set by selecting those with no significant change in IR (IR difference < 0.0005, 0.9 < IR FC < 1.1, and p > 0.5) in either SFPQ knockdown conditions. For each gene set, the percentage of genes with introns overlapping SFPQ CLIP-seq peaks was calculated. Furthermore, for each gene we evaluated the SFPQ peak density as the number of SFPQ peaks on the gene divided by the gene length.

### Semi-quantitative and Quantitative PCR

For both experiments, Control, and Wnt1-Cre; Prmt1fl/fl mandibles, and siRNA-transfected CNCCs, mRNAs were quantified by real-time PCR with IQ Sybr Green Supermix (Bio-Rad) and normalized against GAPDH mRNA levels. Relative changes in expression were calculated using the ΔΔCt method. Primer sequences are listed in Supplemental Table S9.

### Cleavage Under Target and Tagmentation (CUT&Tag)

CUT&Tag was adapted from in Epicypher DIY protocol and Kaya-Okur, H.S., Janssens, D.H., Henikoff, J.G. *et al*. Efficient low-cost chromatin profiling with CUT&Tag. *Nat Protoc* **15**, 3264–3283 (2020). https://doi.org/10.1038/s41596-020-0373-x. Before extracted nuclei, 10 µL of ConA beads was resuspend twice in Bead Activation Buffer (20mM HEPES, pH7.9; 10mM KCl; 1mM CaCl_2_; 1mM MnCl_2_). 100,000 cells were washed once with PBS and and nuclei were extracted by resuspending cells in nuclear extraction (NE) buffer (20mM HEPES-KOH, pH 7.9, 20mM KCl, 0.1% Triton X-100, 20% Glycerol, 0.5mM Spermidine, and protease inhibitor) and incubating for 10 minutes on ice. Nuclei were collected by centrifuge at 600g for 3 minutes and resuspended in 100 µL of NE buffer per 100,000 nuclei. 100 µL of nuclei were transferred to PCR tube containing 10 µL of activated ConA beads, gently vortex, and incubate at room temp for 10 minutes. The supernatant was discarded and 50 µL of Antibody buffer (20mM HEPES, pH7.5; 150mM NaCl; 0.5mM Spermidine; 0.01% Digitonin; 2mM ETDA; and protease inhibitor) was added. An antibody against RNA Pol subunit 1 (Rbp1 CTD (4H8) mouse mAbm Cell Signal Tech.) was added and incubated on a nutator at 4C overnight. Beads was then incubated in 50µL of 10 µg/ml of secondary antibody was made in Digitonin150 buffer (20mM HEPES, pH7.5; 150mM NaCl; 0.5mM Spermidine; 0.01% Digitonin; and protease inhibitor) at room temperature for 30 minutes on a nutator. Wash twice with Digitonin150 buffer before resuspending beads in pAG-Tn5 diluted in Digitonin300 buffer (20mM HEPES, pH7.5; 300mM NaCl; 0.5mM Spermidine; 0.01% Digitonin; and protease inhibitor) and incubating for 1 hour at room temperature on a nutator. Wash twice with Digitonin300 buffer then start the tagmentation reaction by resuspending beads in Tagmentation buffer (Digitonin300 supplemented with 10mM MgCl_2_) and incubated at 37C in thermocycler. Beads were washed once with 50µL TAPS Buffer (10mM TAPS pH 8.5; 0.2mM EDTA) then incubated in 50µL of Release Buffer (10mM TAPS pH 8.5; 0.1% SDS; 0.4 mg/µL Proteinase K; 15mM EDTA) at 55C for 1 hour; 75C for 20 minutes; and then cool down to 20C. Tagmented DNA were purified with 1.8x AMPureXP beads and eluted with 20µL of DNase/RNase free H_2_O. The library was prepared with dual index primers and NEBNext High-Fidelity 2X PCR master mix (M0541) and amplified for 14 cycles. Library DNA was purified with 1.3x AMPureXP beads and eluted with 15µL 0.1xTE buffer. Sequencing run was performed on the NovaSeq X Series platform with 10B reagent kit (300 Cycle). NovaSeq Control Software 1.2.2.48004 was used for sequencing and Illumina BCL Convert v4.2.7 to convert base call (BCL) files into FASTQ files.

### Statistical analysis

Two-tailed Student’s t-tests or Fisher’s exact tests were applied for statistical analysis. For all graphs, error bars represent standard deviations. A P-value of <0.05 was considered statistically significant.

## Acknowledgement

We thank the Flow Cytometry Facility of USC Stem Cell at the University of Southern California. This research was supported by NIH NIDCR R01DE028943 (to J.X.), NIDCR U01DE022937 (to S.Y.) and NIAM R00AR077090 (to Z.L.), NIDCR T90 grant (to T.R.), and Anandamahidol Foundation Scholarship (to. N.U.).

## Author contributions

J.R. and N.U. designed and performed experiments, analyzed data, drafted, and revised the manuscript; Q.C. and W.P. performed bioinformatic analysis of RNA-seq data and conducted intron analysis; M.V., Z.L., S.Y., Y.C. and A.M. contributed to embryo collection and CNCC isolation; T.R. contributed to RT-PCR analysis, H.P. and Y.Y. performed the Pol II CUT&RUN, J.X., and W.P. concepted the study, supervised the performance of experiments and data analysis, and critically revised the manuscript.

## Data Availability

RNA-seq data is deposited at the NCBI Gene Expression Omnibus (GSE 171630)

## REFERENCES

1. Achilleos, A., & Trainor, P. A. (2012). Neural crest stem cells: discovery, properties and potential for therapy. Cell Res, 22(2), 288–304. doi:10.1038/cr.2012.11

2. Anders, S., Pyl, P. T., & Huber, W. (2015). HTSeq--a Python framework to work with high-throughput sequencing data. Bioinformatics, 31(2), 166–169. doi:10.1093/bioinformatics/btu638

3. Bélanger, C., Bérubé-Simard, F. A., Leduc, E., Bernas, G., Campeau, P. M., Lalani, S. R., . . . Pilon, N. (2018). Dysregulation of cotranscriptional alternative splicing underlies CHARGE syndrome. Proc Natl Acad Sci U S A, 115(4), E620–e629. doi:10.1073/pnas.1715378115

4. Berger, H., Wodarz, A., & Borchers, A. (2017). PTK7 Faces the Wnt in Development and Disease. Front Cell Dev Biol, 5, 31. doi:10.3389/fcell.2017.00031

5. Blanc, R. S., & Richard, S. (2017). Arginine Methylation: The Coming of Age. Mol Cell, 65(1), 8–24. doi:10.1016/j.molcel.2016.11.003

6. Braunschweig, U., Barbosa-Morais, N. L., Pan, Q., Nachman, E. N., Alipanahi, B., Gonatopoulos-Pournatzis, T., . . . Blencowe, B. J. (2014). Widespread intron retention in mammals functionally tunes transcriptomes. Genome Res, 24(11), 1774–1786. doi:10.1101/gr.177790.114

7. Breschi, A., Gingeras, T. R., & Guigó, R. (2017). Comparative transcriptomics in human and mouse. Nat Rev Genet, 18(7), 425–440. doi:10.1038/nrg.2017.19

8. Canales, D., Moyano, D., Alvarez, F., Grande-Tovar, C. D., Valencia-Llano, C. H., Peponi, L., . . . Zapata, P. A. (2023). Preparation and characterization of novel poly (lactic acid)/calcium oxide nanocomposites by electrospinning as a potential bone tissue scaffold. Biomater Adv, 153, 213578. doi:10.1016/j.bioadv.2023.213578

9. Chai, Y., Jiang, X., Ito, Y., Bringas, P., Jr., Han, J., Rowitch, D. H., . . . Sucov, H. M. (2000). Fate of the mammalian cranial neural crest during tooth and mandibular morphogenesis. Development, 127(8), 1671–1679. doi:10.1242/dev.127.8.1671

10. Chen, L., Zhang, M., Fang, L., Yang, X., Cao, N., Xu, L., . . . Cao, Y. (2021). Coordinated regulation of the ribosome and proteasome by PRMT1 in the maintenance of neural stemness in cancer cells and neural stem cells. J Biol Chem, 297(5), 101275. doi:10.1016/j.jbc.2021.101275

11. Cibi, D. M., Mia, M. M., Guna Shekeran, S., Yun, L. S., Sandireddy, R., Gupta, P., . . . Singh, M. K. (2019). Neural crest-specific deletion of Rbfox2 in mice leads to craniofacial abnormalities including cleft palate. Elife, 8. doi:10.7554/eLife.45418

12. Cosker, K. E., Fenstermacher, S. J., Pazyra-Murphy, M. F., Elliott, H. L., & Segal, R. A. (2016). The RNA-binding protein SFPQ orchestrates an RNA regulon to promote axon viability. Nat Neurosci, 19(5), 690–696. doi:10.1038/nn.4280

13. Dvinge, H., & Bradley, R. K. (2015). Widespread intron retention diversifies most cancer transcriptomes. Genome Med, 7(1), 45. doi:10.1186/s13073-015-0168-9

14. Fedoriw, A., Rajapurkar, S. R., O’Brien, S., Gerhart, S. V., Mitchell, L. H., Adams, N. D., . . . Mohammad, H. P. (2019). Anti-tumor Activity of the Type I PRMT Inhibitor, GSK3368715, Synergizes with PRMT5 Inhibition through MTAP Loss. Cancer Cell, 36(1), 100–114.e125. doi:10.1016/j.ccell.2019.05.014

15. Feng, Z., Duren, Z., Xiong, Z., Wang, S., Liu, F., Wong, W. H., & Wang, Y. (2021). hReg-CNCC reconstructs a regulatory network in human cranial neural crest cells and annotates variants in a developmental context. Commun Biol, 4(1), 442. doi:10.1038/s42003-021-01970-0

16. Fong, J. Y., Pignata, L., Goy, P. A., Kawabata, K. C., Lee, S. C., Koh, C. M., . . . Guccione, E. (2019). Therapeutic Targeting of RNA Splicing Catalysis through Inhibition of Protein Arginine Methylation. Cancer Cell, 36(2), 194–209.e199. doi:10.1016/j.ccell.2019.07.003

17. Glessner, J. T., Bick, A. G., Ito, K., Homsy, J., Rodriguez-Murillo, L., Fromer, M., . . . Chung, W. K. (2014). Increased frequency of de novo copy number variants in congenital heart disease by integrative analysis of single nucleotide polymorphism array and exome sequence data. Circ Res, 115(10), 884–896. doi:10.1161/circresaha.115.304458

18. Gou, Y., Li, J., Jackson-Weaver, O., Wu, J., Zhang, T., Gupta, R., . . . Xu, J. (2018). Protein Arginine Methyltransferase PRMT1 Is Essential for Palatogenesis. J Dent Res, 97(13), 1510–1518.

19. Gou, Y., Li, J., Wu, J., Gupta, R., Cho, I., Ho, T. V., . . . Xu, J. (2018). Prmt1 regulates craniofacial bone formation upstream of Msx1. Mech Dev, 152, 13–20. doi:10.1016/j.mod.2018.05.001

20. Griffin, C., & Saint-Jeannet, J. P. (2020). Spliceosomopathies: Diseases and mechanisms. Dev Dyn, 249(9), 1038–1046.

21. Han, X., Wei, Y., Wang, H., Wang, F., Ju, Z., & Li, T. (2018). Nonsense-mediated mRNA decay: a ’nonsense’ pathway makes sense in stem cell biology. Nucleic Acids Res, 46(3), 1038–1051. doi:10.1093/nar/gkx1272

22. Hartel, N. G., Chew, B., Qin, J., Xu, J., & Graham, N. A. (2019). Deep Protein Methylation Profiling by Combined Chemical and Immunoaffinity Approaches Reveals Novel PRMT1 Targets. Mol Cell Proteomics, 18(11), 2149–2164. doi:10.1074/mcp.RA119.001625

23. Hashimoto, M., Fukamizu, A., Nakagawa, T., & Kizuka, Y. (2021). Roles of protein arginine methyltransferase 1 (PRMT1) in brain development and disease. Biochim Biophys Acta Gen Subj, 1865(1), 129776. doi:10.1016/j.bbagen.2020.129776

24. Hashimoto, M., Kumabe, A., Kim, J. D., Murata, K., Sekizar, S., Williams, A., . . . Fukamizu, A. (2021). Loss of PRMT1 in the central nervous system (CNS) induces reactive astrocytes and microglia during postnatal brain development. J Neurochem, 156(6), 834–847. doi:10.1111/jnc.15149

25. Hashimoto, M., Takeichi, K., Murata, K., Kozakai, A., Yagi, A., Ishikawa, K., . . . Nakagawa, T. (2022). Regulation of neural stem cell proliferation and survival by protein arginine methyltransferase 1. Front Neurosci, 16, 948517. doi:10.3389/fnins.2022.948517

26. Homsy, J., Zaidi, S., Shen, Y., Ware, J. S., Samocha, K. E., Karczewski, K. J., . . . Chung, W. K. (2015). De novo mutations in congenital heart disease with neurodevelopmental and other congenital anomalies. Science, 350(6265), 1262–1266. doi:10.1126/science.aac9396

27. Hong, X., Scofield, D. G., & Lynch, M. (2006). Intron size, abundance, and distribution within untranslated regions of genes. Mol Biol Evol, 23(12), 2392–2404. doi:10.1093/molbev/msl111

28. Hooper, J. E., Jones, K. L., Smith, F. J., Williams, T., & Li, H. (2020). An Alternative Splicing Program for Mouse Craniofacial Development. Front Physiol, 11, 1099. doi:10.3389/fphys.2020.01099

29. Hosokawa, M., Takeuchi, A., Tanihata, J., Iida, K., Takeda, S., & Hagiwara, M. (2019). Loss of RNA-Binding Protein Sfpq Causes Long-Gene Transcriptopathy in Skeletal Muscle and Severe Muscle Mass Reduction with Metabolic Myopathy. iScience, 13, 229–242.

30. Hwang, J. Y., Jung, S., Kook, T. L., Rouchka, E. C., Bok, J., & Park, J. W. (2020). rMAPS2: an update of the RNA map analysis and plotting server for alternative splicing regulation. Nucleic Acids Res, 48(W1), W300–w306. doi:10.1093/nar/gkaa237

31. Ishida, M., Kawao, N., Mizukami, Y., Takafuji, Y., & Kaji, H. (2021). Influence of Angptl1 on osteoclast formation and osteoblastic phenotype in mouse cells. BMC Musculoskelet Disord, 22(1), 398. doi:10.1186/s12891-021-04278-6

32. Ishihara, S., Usumi-Fujita, R., Kasahara, Y., Oishi, S., Shibata, K., Shimizu, Y., . . . Ono, T. (2023). Periostin splice variants affect craniofacial growth by influencing chondrocyte hypertrophy. J Bone Miner Metab, 41(2), 171–181. doi:10.1007/s00774-023-01409-y

33. Ishii, M., Arias, A. C., Liu, L., Chen, Y. B., Bronner, M. E., & Maxson, R. E. (2012). A stable cranial neural crest cell line from mouse. Stem Cells Dev, 21(17), 3069–3080. doi:10.1089/scd.2012.0155

34. Jackson-Weaver, O., Ungvijanpunya, N., Yuan, Y., Qian, J., Gou, Y., Wu, J., . . . Xu, J. (2020). PRMT1-p53 Pathway Controls Epicardial EMT and Invasion. Cell Rep, 31(10), 107739.

35. Jiang, X., Iseki, S., Maxson, R. E., Sucov, H. M., & Morriss-Kay, G. M. (2002). Tissue origins and interactions in the mammalian skull vault. Dev Biol, 241(1), 106–116. doi:10.1006/dbio.2001.0487

36. Jobert, L., Argentini, M., & Tora, L. (2009). PRMT1 mediated methylation of TAF15 is required for its positive gene regulatory function. Exp Cell Res, 315(7), 1273–1286. doi:10.1016/j.yexcr.2008.12.008

37. Kim, D., Paggi, J. M., Park, C., Bennett, C., & Salzberg, S. L. (2019). Graph-based genome alignment and genotyping with HISAT2 and HISAT-genotype. Nat Biotechnol, 37(8), 907–915. doi:10.1038/s41587-019-0201-4

38. Li, H., Handsaker, B., Wysoker, A., Fennell, T., Ruan, J., Homer, N., . . . Durbin, R. (2009). The Sequence Alignment/Map format and SAMtools. Bioinformatics, 25(16), 2078–2079. doi:10.1093/bioinformatics/btp352

39. Li, K., Xiu, C., Zhou, Q., Ni, L., Du, J., Gong, T., . . . Chen, J. (2019). A dual role of cholesterol in osteogenic differentiation of bone marrow stromal cells. J Cell Physiol, 234(3), 2058–2066. doi:10.1002/jcp.27635

40. Li, K. K. C., Chau, B. L., & Lee, K. A. W. (2018). Differential interaction of PRMT1 with RGG-boxes of the FET family proteins EWS and TAF15. Protein Sci, 27(3), 633–642. doi:10.1002/pro.3354

41. Liu, Q., & Dreyfuss, G. (1995). In vivo and in vitro arginine methylation of RNA-binding proteins. Mol Cell Biol, 15(5), 2800–2808. doi:10.1128/MCB.15.5.2800

42. Lopes, I., Altab, G., Raina, P., & de Magalhães, J. P. (2021). Gene Size Matters: An Analysis of Gene Length in the Human Genome. Front Genet, 12, 559998. doi:10.3389/fgene.2021.559998

43. Luisier, R., Tyzack, G. E., Hall, C. E., Mitchell, J. S., Devine, H., Taha, D. M., . . . Patani, R. (2018). Intron retention and nuclear loss of SFPQ are molecular hallmarks of ALS. Nat Commun, 9(1), 2010. doi:10.1038/s41467-018-04373-8

44. Marquez, Y., Brown, J. W., Simpson, C., Barta, A., & Kalyna, M. (2012). Transcriptome survey reveals increased complexity of the alternative splicing landscape in Arabidopsis. Genome Res, 22(6), 1184–1195. doi:10.1101/gr.134106.111

45. Martik, M. L., & Bronner, M. E. (2017). Regulatory Logic Underlying Diversification of the Neural Crest. Trends Genet, 33(10), 715–727. doi:10.1016/j.tig.2017.07.015

46. Martin, L., Grigoryan, A., Wang, D., Wang, J., Breda, L., Rivella, S., . . . Gardner, L. B. (2014). Identification and characterization of small molecules that inhibit nonsense-mediated RNA decay and suppress nonsense p53 mutations. Cancer Res, 74(11), 3104–3113.

47. Mauger, O., Lemoine, F., & Scheiffele, P. (2016). Targeted Intron Retention and Excision for Rapid Gene Regulation in Response to Neuronal Activity. Neuron, 92(6), 1266–1278. doi:10.1016/j.neuron.2016.11.032

48. McKean, D. M., Homsy, J., Wakimoto, H., Patel, N., Gorham, J., DePalma, S. R., . . . Seidman, J. G. (2016). Loss of RNA expression and allele-specific expression associated with congenital heart disease. Nat Commun, 7, 12824. doi:10.1038/ncomms12824

49. Merkuri, F., & Fish, J. L. (2019). Developmental processes regulate craniofacial variation in disease and evolution. Genesis, 57(1), e23249. doi:10.1002/dvdy.214

50. Mizukami, Y., Kawao, N., Takafuji, Y., Ohira, T., Okada, K., Jo, J. I., . . . Kaji, H. (2023). Matrix vesicles promote bone repair after a femoral bone defect in mice. PLoS One, 18(4), e0284258. doi:10.1371/journal.pone.0284258

51. Mizumoto, S., & Yamada, S. (2021). Congenital Disorders of Deficiency in Glycosaminoglycan Biosynthesis. Front Genet, 12, 717535. doi:10.3389/fgene.2021.717535

52. Monteuuis, G., Wong, J. J. L., Bailey, C. G., Schmitz, U., & Rasko, J. E. J. (2019). The changing paradigm of intron retention: regulation, ramifications and recipes. Nucleic Acids Res, 47(22), 11497–11513. doi:10.1093/nar/gkz1068

53. Ni, T., Yang, W., Han, M., Zhang, Y., Shen, T., Nie, H., . . . Zhu, J. (2016). Global intron retention mediated gene regulation during CD4+ T cell activation. Nucleic Acids Res, 44(14), 6817–6829. doi:10.1093/nar/gkw591

54. Paganini, C., Gramegna Tota, C., Superti-Furga, A., & Rossi, A. (2020). Skeletal Dysplasias Caused by Sulfation Defects. Int J Mol Sci, 21(8). doi:10.3390/ijms21082710

55. Pahlich, S., Bschir, K., Chiavi, C., Belyanskaya, L., & Gehring, H. (2005). Different methylation characteristics of protein arginine methyltransferase 1 and 3 toward the Ewing Sarcoma protein and a peptide. Proteins, 61(1), 164–175. doi:10.1002/prot.20579

56. Park, J. W., Jung, S., Rouchka, E. C., Tseng, Y. T., & Xing, Y. (2016). rMAPS: RNA map analysis and plotting server for alternative exon regulation. Nucleic Acids Res, 44(W1), W333–338. doi:10.1093/nar/gkw410

57. Plein, A., Fantin, A., & Ruhrberg, C. (2015). Neural crest cells in cardiovascular development. Curr Top Dev Biol, 111, 183–200. doi:10.1016/bs.ctdb.2014.11.006

58. Pregizer, S., Barski, A., Gersbach, C. A., García, A. J., & Frenkel, B. (2007). Identification of novel Runx2 targets in osteoblasts: cell type-specific BMP-dependent regulation of Tram2. J Cell Biochem, 102(6), 1458–1471. doi:10.1002/jcb.21366

59. Rho, J., Choi, S., Jung, C. R., & Im, D. S. (2007). Arginine methylation of Sam68 and SLM proteins negatively regulates their poly(U) RNA binding activity. Arch Biochem Biophys, 466(1), 49–57. doi:10.1016/j.abb.2007.07.017

60. Robinson, M. D., McCarthy, D. J., & Smyth, G. K. (2010). edgeR: a Bioconductor package for differential expression analysis of digital gene expression data. Bioinformatics, 26(1), 139–140. doi:10.1093/bioinformatics/btp616

61. Ruskin, B., Zamore, P. D., & Green, M. R. (1988). A factor, U2AF, is required for U2 snRNP binding and splicing complex assembly. Cell, 52(2), 207–219. doi:10.1016/0092-8674(88)90509-0

62. Ruta, V., Pagliarini, V., & Sette, C. (2021). Coordination of RNA Processing Regulation by Signal Transduction Pathways. Biomolecules, 11(10). doi:10.3390/biom11101475

63. Schmitz, U., Pinello, N., Jia, F., Alasmari, S., Ritchie, W., Keightley, M. C., . . . Rasko, J. E. J. (2017). Intron retention enhances gene regulatory complexity in vertebrates. Genome Biol, 18(1), 216.

64. Schwartz, N. B., & Domowicz, M. (2002). Chondrodysplasias due to proteoglycan defects. Glycobiology, 12(4), 57r–68r. doi:10.1093/glycob/12.4.57r

65. Seong, C. H., Chiba, N., Fredy, M., Kusuyama, J., Ishihata, K., Kibe, T., . . . Matsuguchi, T. (2023). Early induction of Hes1 by bone morphogenetic protein 9 plays a regulatory role in osteoblastic differentiation of a mesenchymal stem cell line. J Cell Biochem, 124(9), 1366–1378. doi:10.1002/jcb.30452

66. Shaheen, R., Anazi, S., Ben-Omran, T., Seidahmed, M. Z., Caddle, L. B., Palmer, K., . . . Alkuraya, F. S. (2016). Mutations in SMG9, Encoding an Essential Component of Nonsense-Mediated Decay Machinery, Cause a Multiple Congenital Anomaly Syndrome in Humans and Mice. Am J Hum Genet, 98(4), 643–652. doi:10.1016/j.ajhg.2016.02.010

67. Sherman, B. T., Hao, M., Qiu, J., Jiao, X., Baseler, M. W., Lane, H. C., . . . Chang, W. (2022). DAVID: a web server for functional enrichment analysis and functional annotation of gene lists (2021 update). Nucleic Acids Res, 50(W1), W216–w221. doi:10.1093/nar/gkac194

68. Smith, W. A., Schurter, B. T., Wong-Staal, F., & David, M. (2004). Arginine methylation of RNA helicase a determines its subcellular localization. J Biol Chem, 279(22), 22795–22798. doi:10.1074/jbc.C300512200

69. Snijders, A. P., Hautbergue, G. M., Bloom, A., Williamson, J. C., Minshull, T. C., Phillips, H. L., . . . Dickman, M. J. (2015). Arginine methylation and citrullination of splicing factor proline- and glutamine-rich (SFPQ/PSF) regulates its association with mRNA. Rna, 21(3), 347–359. doi:10.1261/rna.045138.114

70. Strauss, F. J., Di Summa, F., Stähli, A., Matos, L., Vaca, F., Schuldt, G., & Gruber, R. (2019). TGF-β activity in acid bone lysate adsorbs to titanium surface. Clin Implant Dent Relat Res, 21(2), 336–343. doi:10.1111/cid.12734

71. Strobl-Mazzulla, P. H., Marini, M., & Buzzi, A. (2012). Epigenetic landscape and miRNA involvement during neural crest development. Dev Dyn, 241(12), 1849–1856. doi:10.1002/dvdy.23868

72. Sun, X., Liu, Z., Li, Z., Zeng, Z., Peng, W., Zhu, J., . . . Lipsky, P.E. (2023). Abnormalities in intron retention characterize patients with systemic lupus erythematosus. Sci Rep, 13(1), 5141. doi:10.1038/s41598-023-31890-4

73. Sun, X., Moriarty, P. M., & Maquat, L. E. (2000). Nonsense-mediated decay of glutathione peroxidase 1 mRNA in the cytoplasm depends on intron position. EMBO J, 19(17), 4734–4744. doi:10.1093/emboj/19.17.4734

74. Takeuchi, A., Iida, K., Tsubota, T., Hosokawa, M., Denawa, M., Brown, J. B., . . . Hagiwara, M. (2018). Loss of Sfpq Causes Long-Gene Transcriptopathy in the Brain. Cell Rep, 23(5), 1326–1341. doi:10.1016/j.celrep.2018.03.141

75. Tan, K., Stupack, D. G., & Wilkinson, M. F. (2022). Nonsense-mediated RNA decay: an emerging modulator of malignancy. Nat Rev Cancer, 22(8), 437–451. doi:10.1038/s41568-022-00481-2

76. Taylor, R., Hamid, F., Fielding, T., Gordon, P. M., Maloney, M., Makeyev, E. V., & Houart, C. (2022). Prematurely terminated intron-retaining mRNAs invade axons in SFPQ null-driven neurodegeneration and are a hallmark of ALS. Nat Commun, 13(1), 6994. doi:10.1038/s41467-022-34331-4

77. Thandapani, P., O’Connor, T. R., Bailey, T. L., & Richard, S. (2013). Defining the RGG/RG motif. Mol Cell, 50(5), 613–623. doi:10.1016/j.molcel.2013.05.021

78. Thomas-Jinu, S., Gordon, P. M., Fielding, T., Taylor, R., Smith, B. N., Snowden, V., . . . Houart, C. (2017). Non-nuclear Pool of Splicing Factor SFPQ Regulates Axonal Transcripts Required for Normal Motor Development. Neuron, 94(2), 322–336.e325. doi:10.1016/j.neuron.2017.03.026

79. Tian, Y., Zeng, Z., Li, X., Wang, Y., Chen, R., Mattijssen, S., . . . Zhu, J. (2020). Transcriptome-wide stability analysis uncovers LARP4-mediated NFκB1 mRNA stabilization during T cell activation. Nucleic Acids Res, 48(15), 8724–8739. doi:10.1093/nar/gkaa643

80. Tu, X., Joeng, K. S., Nakayama, K. I., Nakayama, K., Rajagopal, J., Carroll, T. J., . . . Long, F. (2007). Noncanonical Wnt signaling through G protein-linked PKCdelta activation promotes bone formation. Dev Cell, 12(1), 113–127. doi:10.1016/j.devcel.2006.11.003

81. Verma, S. K., Deshmukh, V., Thatcher, K., Belanger, K. K., Rhyner, A. M., Meng, S., . . . Kuyumcu-Martinez, M. N. (2022). RBFOX2 is required for establishing RNA regulatory networks essential for heart development. Nucleic Acids Res, 50(4), 2270–2286. doi:10.1093/nar/gkac055

82. Wang, E. T., Sandberg, R., Luo, S., Khrebtukova, I., Zhang, L., Mayr, C., . . . Burge, C. B. (2008). Alternative isoform regulation in human tissue transcriptomes. Nature, 456(7221), 470–476. doi:10.1038/nature07509

83. Wang, Y., Xie, Z., Kutschera, E., Adams, J. I., Kadash-Edmondson, K. E., & Xing, Y. (2024). rMATS-turbo: an efficient and flexible computational tool for alternative splicing analysis of large-scale RNA-seq data. Nat Protoc, 19(4), 1083–1104. doi:10.1038/s41596-023-00944-2

84. Wong, J. J., Au, A. Y., Ritchie, W., & Rasko, J. E. (2016). Intron retention in mRNA: No longer nonsense: Known and putative roles of intron retention in normal and disease biology. Bioessays, 38(1), 41–49. doi:10.1002/bies.201500117

85. Wong, J. J., & Schmitz, U. (2022). Intron retention: importance, challenges, and opportunities. Trends Genet, 38(8), 789–792. doi:10.1016/j.tig.2022.03.017

86. Wu, H., Zhang, Y., Liu, S., Liu, D., Li, A., Deng, H., . . . Pang, Q. (2022). Protein Arginine Methyltransferase 1 and its Dynamic Regulation Associated with Cellular Processes and Diseases. Protein Pept Lett, 29(3), 218–230. doi:10.2174/0929866529666220124120208

87. Xu, J., & Richard, S. (2021). Cellular pathways influenced by protein arginine methylation: Implications for cancer. Mol Cell, 81(21), 4357–4368. doi:10.1016/j.molcel.2021.09.011

88. Yang, X., Wang, G., Wang, Y., Zhou, J., Yuan, H., Li, X., . . . Wang, B. (2019). Histone demethylase KDM7A reciprocally regulates adipogenic and osteogenic differentiation via regulation of C/EBPα and canonical Wnt signalling. J Cell Mol Med, 23(3), 2149–2162. doi:10.1111/jcmm.14126

89. Zhang, T., Wu, J., Ungvijanpunya, N., Jackson-Weaver, O., Gou, Y., Feng, J., . . . Xu, J. (2018). Smad6 Methylation Represses NFκB Activation and Periodontal Inflammation. J Dent Res, 97(7), 810–819.

90. Zhou, Y., Zhou, B., Pache, L., Chang, M., Khodabakhshi, A. H., Tanaseichuk, O., . . . Chanda, S. K. (2019). Metascape provides a biologist-oriented resource for the analysis of systems-level datasets. Nat Commun, 10(1), 1523. doi:10.1038/s41467-019-09234-6

