## Supplemental material for "PRMT1-SFPQ regulates intron retention to control matrix gene expression during craniofacial development"

**Supplemental Figure S1. CNCCs labeled by Tdtomato in mouse embryo.** Sagittal sections of *Wnt1-Cre; R26R*$^{tdTomato}$ whole mouse embryo at E13.5 showing the CNCCs (Red, labeled by Tdtomato) and nuclei (Blue, labeled by DAPI). The boxed fields showed higher magnification of the craniofacial structures including the mesenchyme, where CNCCs are located, and epithelium, labeled by the asterisks. Scale bars = 1.0 mm, 500 μm and 100μm, respectively.

*Supplemental Figure S1* *Lima et al., 2025*

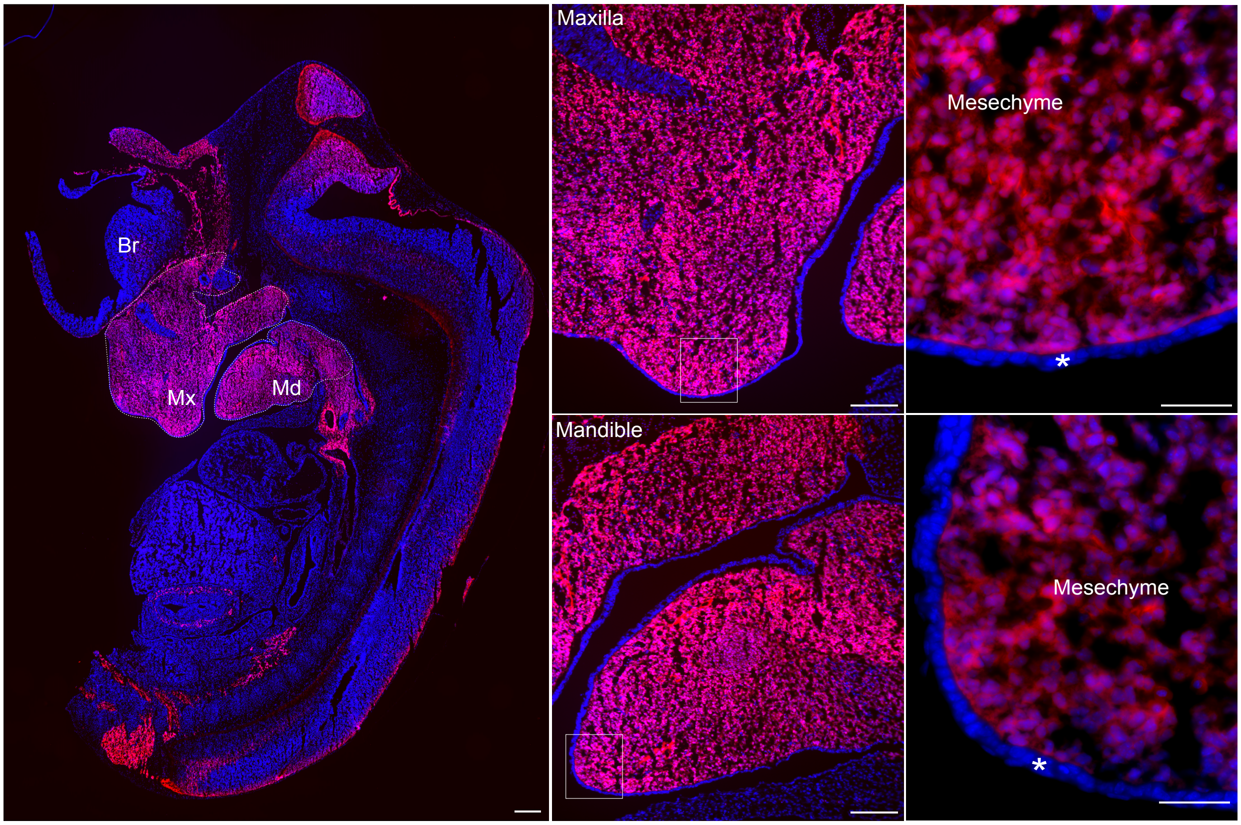


Br: Brain; Mx: Maxilla, Md: Mandible. * Epithelium.

**Supplemental Figure S2. *Prmt1* deletion in CNCCs didn't cause a global shift in intron retention.** Whole genome IRI value distribution in the control (Cont, pink) and *Prmt1* CKO mutant (Mut, blue) embryos was plotted. There is no significant difference between control and mutate intron retention distribution.

*Supplemental Figure S2* *Lima et al., 2025*

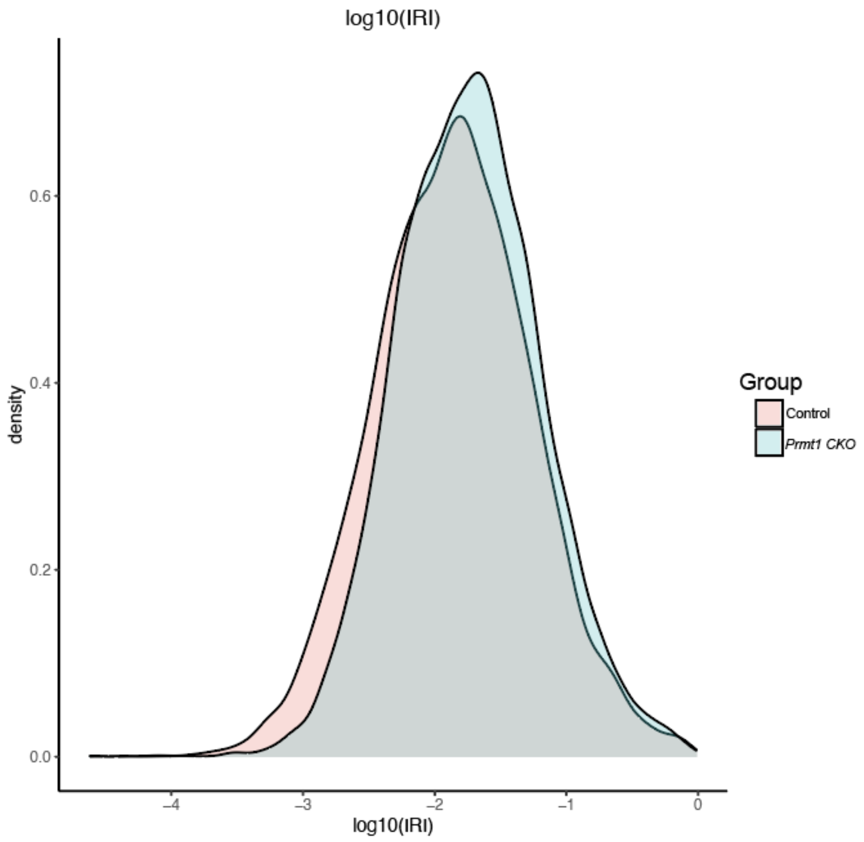

**Supplemental Figure S3. SFPQ, EWSR1, TAF15 and TRA2B methylation in control and *Prmt1 CKO* embryos.** (A) SFPQ methylation signal remained robust in the epithelial region of craniofacial structures in both control (Aa, Ab) and CNCC-specific *Prmt1* deletion (Ac, Ad) embryos. (B) EWSR1 methylation signal was robust in the abdominal region of control (Ba, Bb) and *Prmt1 CKO* (Bc, Bd) embryos. (C) TRA2B methylation signal was robust in the abdominal region of control (Ca, Cb) and *Prmt1 CKO* (Cc, Cd) embryos. Control: *Wnt1-Cre; R26R$^{tdTomato}$*. *Prmt1 CKO: Wnt1-Cre; Prmt1$^{fl/fl}$; R26R$^{tdTomato}$*. Scale bars = 25μm in A, B, C.

*Supplemental Figure S3* *Lima et al., 2025*

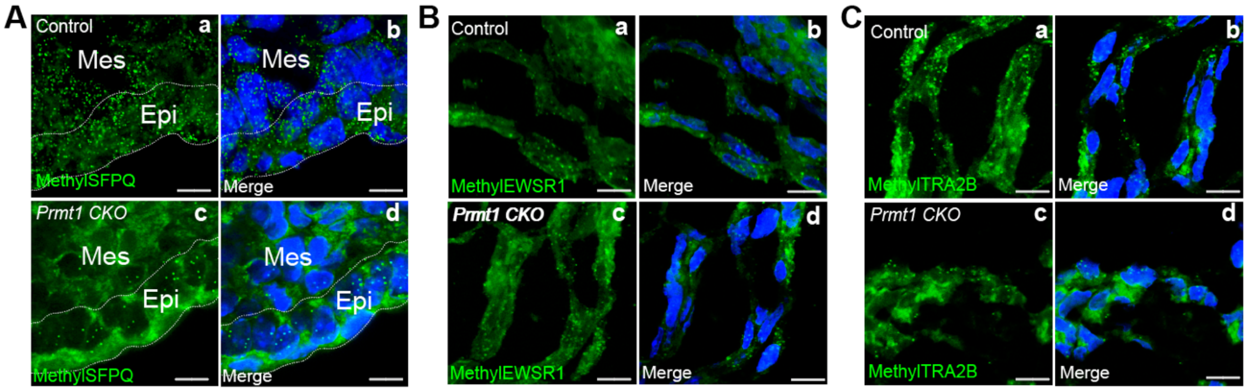

**Supplemental Figure S4. SFPQ, EWSR1, TAF15 and TRA2B protein expression and subcellular localization in control and *Prmt1 CKO* embryos.** EWSR1, TAF15 and TRA2B protein expression and subcellular localization were not altered in the mandibular processes of *Prmt1* deficient embryos. The level of EWSR1 (A), TAF15 (C) and TRA2B (E) protein was detected by immunostaining in the embryonic mandible of control and *Prmt1* deficient embryos. These findings were supported by Western Blot analysis using tissues from the craniofacial structures (G). Subcellular localization was quantified in H-J. Control: *Wnt1-Cre; R26R*$^{tdTomato}$. *Prmt1 CKO: Wnt1-Cre; Prmt1*$^{fl/fl}$*; R26R*$^{tdTomato}$. Scale bars = 100µm in Aa-e, Ca-e, Ea-e. Scale bars = 25µm in Ac, Af, Cc, Cf, Ec, Ef.

*Supplemental Figure S4* *Lima et al., 2025*

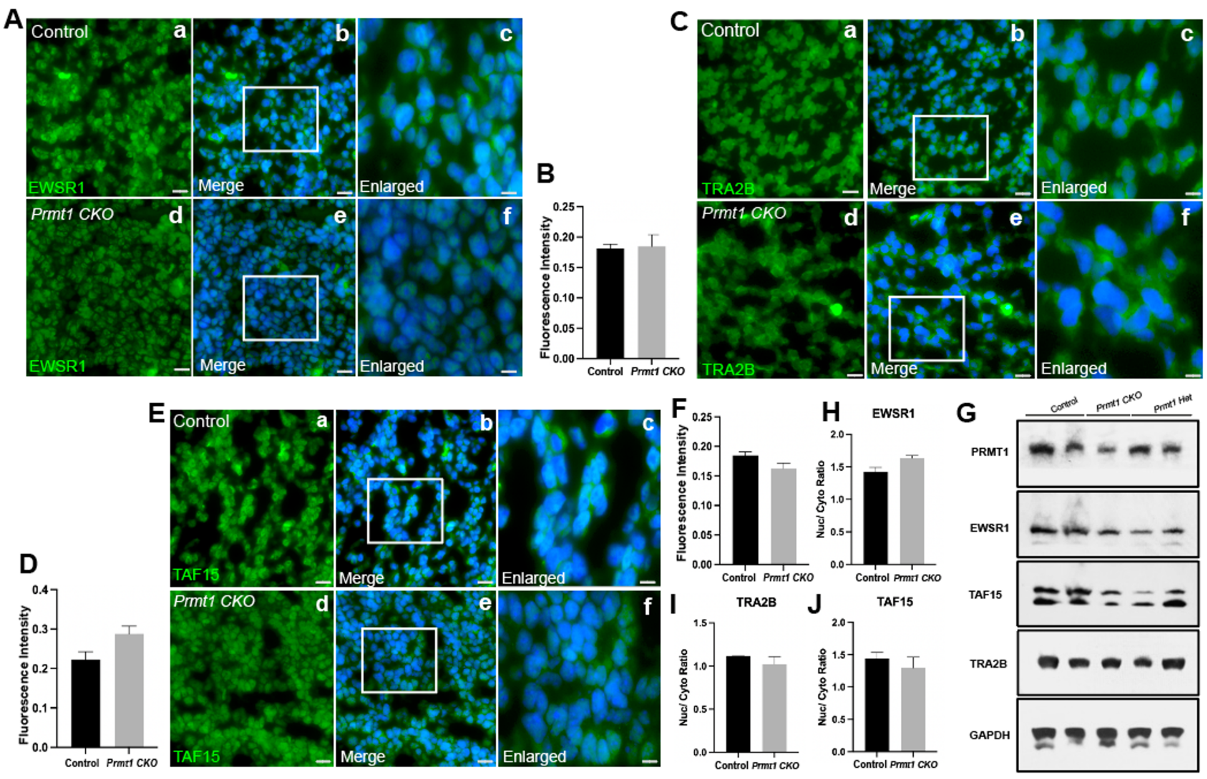

**Supplemental Figure S5. CNCC marker expression in siRNA transfected CNCCs compared to fresh isolated CNCCs from mouse embryos.** Bar chart depicting the CNCC's marker expression in three different conditions. Each condition is presented as mean±SEM.

*Supplemental Figure S5* *Lima et al., 2025*

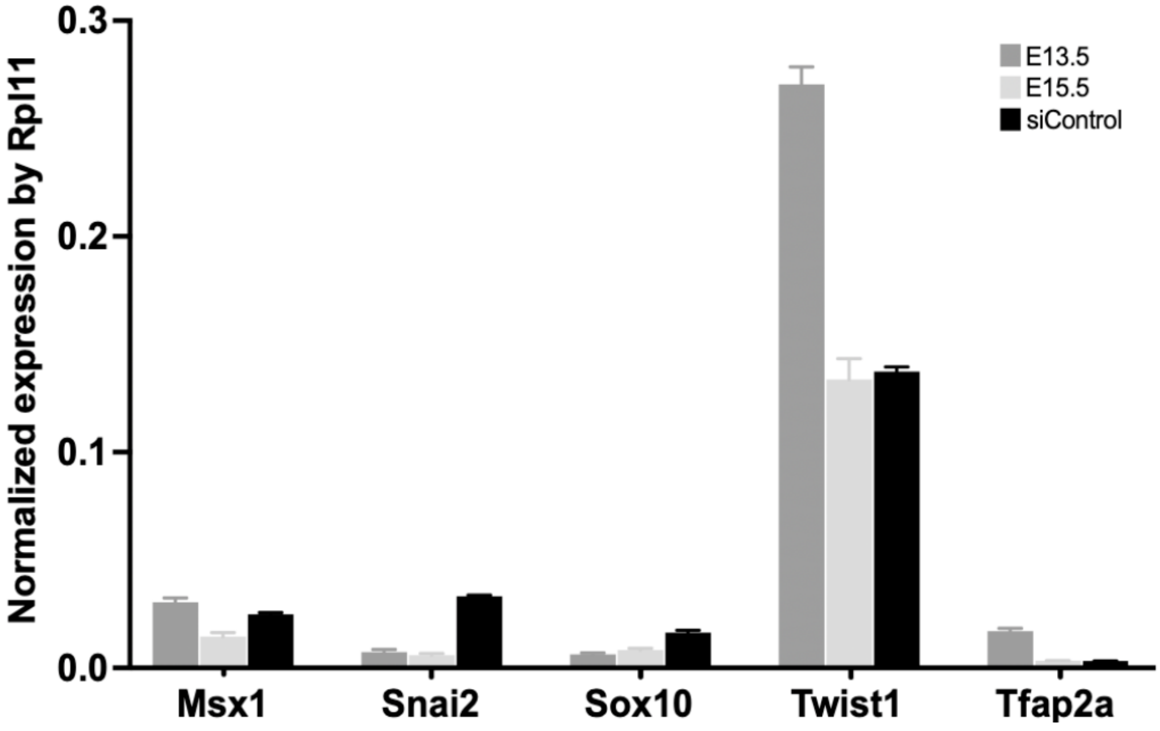

**Supplemental Figure S6. Transcriptional landscape altered by the knockdown of EWSR1, TAF15, and TRA2B.** ST2 cells were transfected with control siRNA or siRNAs targeting EWSR1, TAF15 or TRA2B, followed by mRNA isolation, sequencing, and bioinformatic analysis for GO analysis of upregulated (S6A, S6C, S6E) and downregulated (S6B, S6D, S6F) genes.

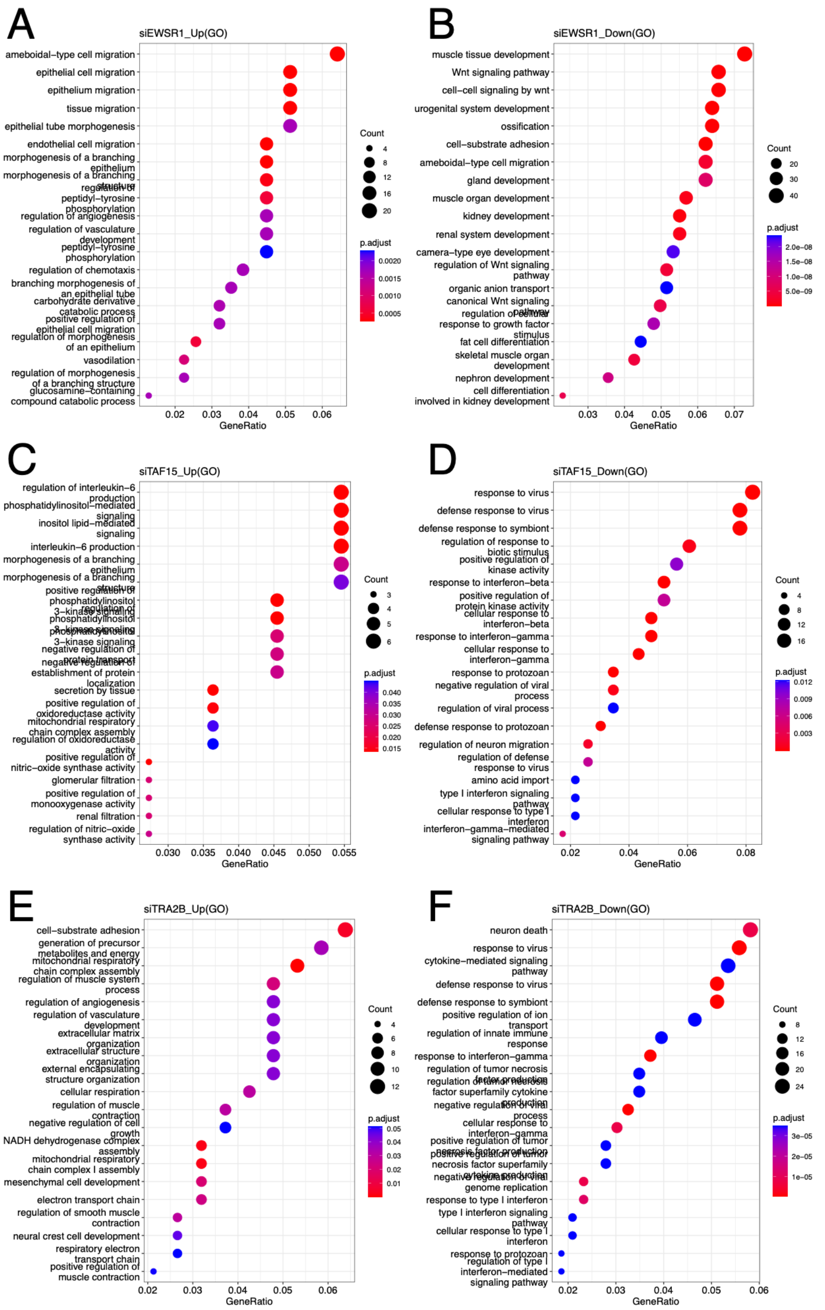

**Supplemental Figure S7. SFPQ motifs in the vicinity of differential alternative splicing events of CNCCs.** rMAPS2 output pages depicting the spatial distribution of SFPQ motifs for Mutually Exclusive Exons (MXE) (A), Exon Skipping (SE) (B), Intron Retention (RI) (C), Alternative 5' Splice Site (A5SS) (D), and Alternative 3' Splice Site (A3SS) (E) events. The results demonstrate high motif scores paired with low P-values for SFPQ, underscoring its significant role in alternative splicing regulation. The red line represents the enriched motif for enhanced exons, the blue line represents the enriched motif for silenced exons, and the black line represents the motif density for background (nonregulated) exons. Solid lines represent the peak quality Motif score (peak height) as scaled on the left. The dotted lines represent the negative log10 (P value) as scaled on the right. The green box indicates the cassette exon.

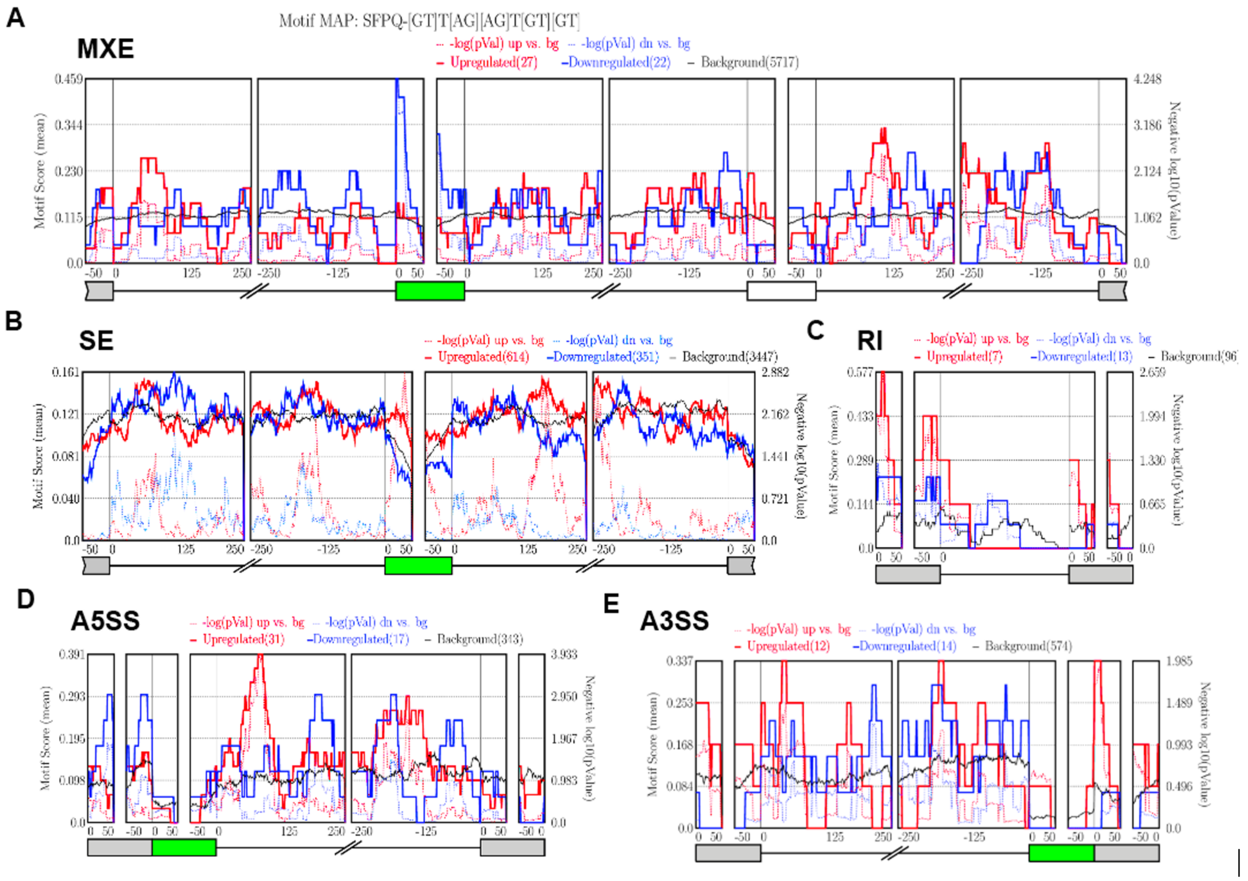


**Supplemental Figure S8. Intron retention coefficient (IRC) analysis provided consistent results as IRI analysis (in response to reviewers' comments).** (A) Intron retention collectively increased for ECM gene transcripts, as illustrated by a scatter plot based on intron retention coefficient (IRC) analysis. This is consistent with IRI analysis shown in Figure 3A. The statistical significance is p-value = 1.3 $10^{-24}$, using one-sided paired-sample Wilcoxon test. (B) Intron retention increased in the majority of GAG degradation gene transcripts that were downregulated in *Prmt1* deficient embryos compared to control, as indicated by a scatter plot based on IRC analysis. This is consistent with IRI analysis shown in Figure 3E. The statistical significance is p-value = 5.5 $10^{-3}$, using one-sided paired-sample Wilcoxon test.

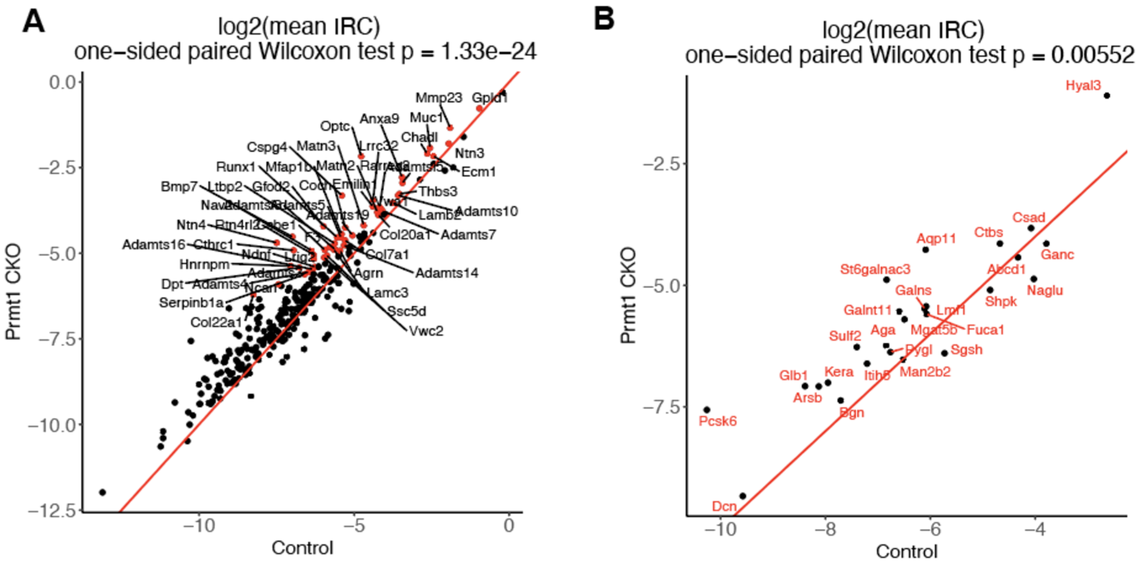


**Supplemental Figure S9. Out of ECM and GAG genes with significant increases in intron retention did not display other types of alternative splicing, reinforcing the role of IR events.** The only ECM and GAG genes with significant non-IR AS changes are illustrated here. (A-E) *Cthcr1*, *Mmp23*, *Adamts10*, *Ccdc80* and *Col25a1* showing statistically significant changes in skipped exon. (F) *Bmp7* showing statistically significant changes in mutually exclusive exons. mean ± SEM. * p<0.05.

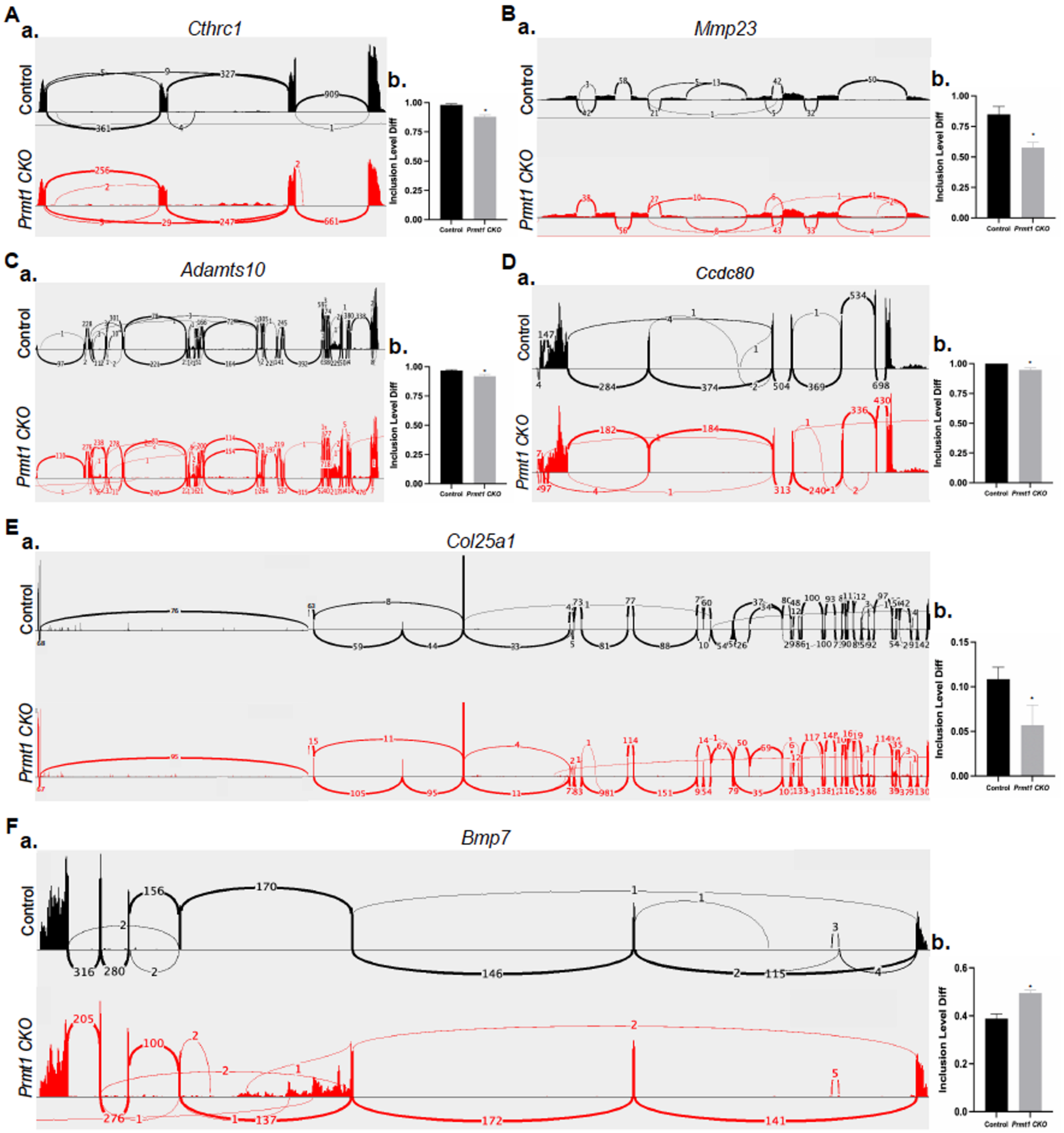


**Supplemental Figure S10. Model illustrating PRMT1-SFPQ-dependent regulation of pre-mRNA splicing and mRNA fate.** In wild-type cells (left panel), PRMT1 methylates the splicing regulator SFPQ, thereby maintaining splicing fidelity, ensuring precise intron removal from pre-mRNA, and generating stable, translation-competent mRNA. In contrast, the loss of PRMT1 or SFPQ activity (right panel) compromises normal splicing, leading to aberrant intron retention within transcripts. These faulty mRNAs are detected by the cellular quality-control pathway Nonsense-Mediated mRNA Decay (NMD) and rapidly degraded.

*Supplemental Figure S10* *Lima et al., 2025*

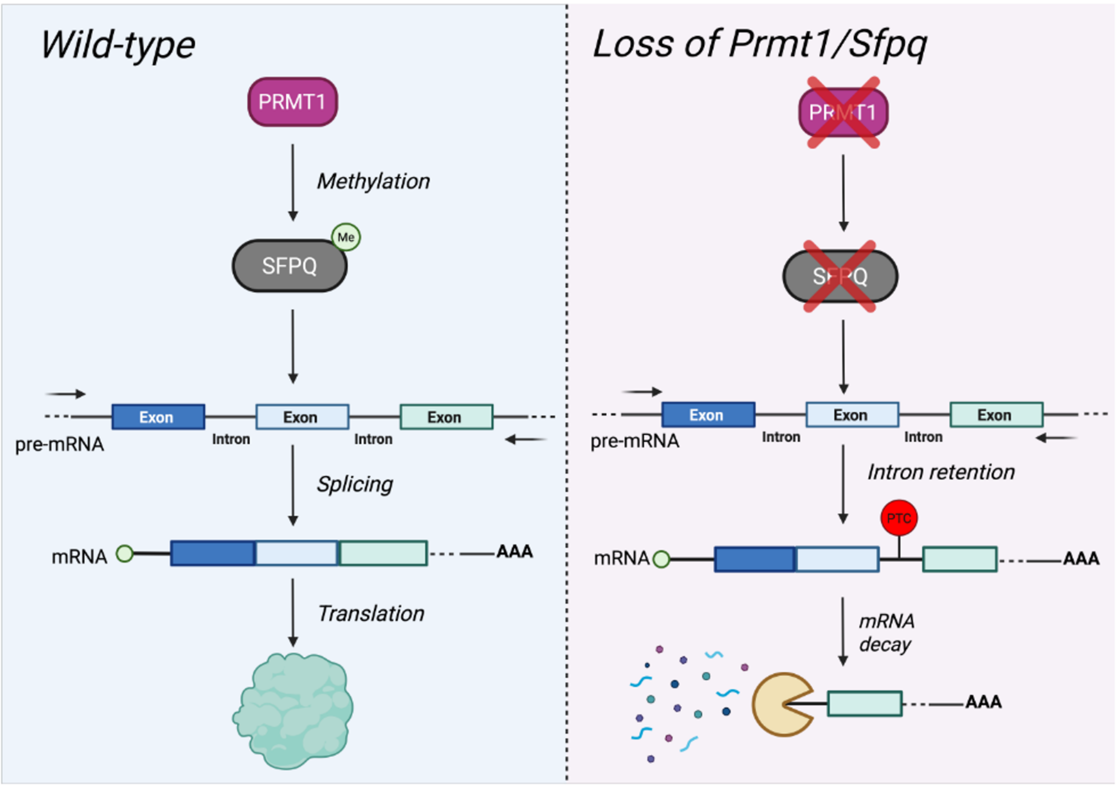


**Supplemental Table S1:** GO analysis of genes with differentially regulated intron retention in *Prmt1 CKO* mandibles**.** Excel file attached.

**Supplemental Table S2:** Number of Premature Termination Codons (PTCs) in the retained introns of ECM and GAG degradation transcripts of *Prmt1 CKO* group.

| Gene | Intron Retention # | Number of Premature Termination Codon (PTC) |
|---|---|---|
| *Bmp7* | 3 | 495 |
| *Muc1* | 1 | 32 |
| *Vwa1* | 1 | 42 |
| *Adamtsl5* | 1 | 13 |
| *Lamb1* | 14 | 52 |
| *Thbs3* | 2 | 34 |
| *Gfod2* | 2 | 401 |
| *Ecm1* | 3 | 4 |
| *Chadl* | 5 | 229 |
| *Plod3* | 18 | 33 |
| *Ltbp3* | 19 | 61 |
| *Anxa9* | 9 | 4 |
| *Emilin1* | 6 | 5 |
| *Serac1* | 12 | 48 |
| *Wnt11* | 3 | 100 |
| *Col7a1* | 100 | 2 |
| *Adamts10* | 9 | 5 |
| *Cthrc1* | 2 | 197 |
| *Cilp* | 1 | 14 |
| *Lingo3* | 1 | 269 |
| *Cspg4* | 4 | 16 |
| *Lamc3* | 11 | 26 |
| *Adamts5* | 6 | 67 |
| *Runx1* | 1 | 282 |
| *Lgalsl* | 1 | 20 |

| | | |
|---|---|---|
| *Adamts16* | 14 | 68 |
| *Serpine1* | 8 | 13 |
| *Adamts4* | 1 | 39 |
| *Gpc2* | 1 | 22 |
| *Hyal3* | 2 | 37 |
| *Ganc* | 5 | 73 |
| *Csad* | 13 | 4 |
| *Naglu* | 1 | 24 |
| *Aga* | 8 | 100 |
| *Sgsh* | 5 | 13 |
| *Abcd 1* | 8 | 11 |
| *Man2b2* | 18 | 19 |
| *Pygl* | 15 | 3 |
| *Fuca1* | 6 | 108 |
| *Shpk* | 4 | 26 |
| *Galnt11* | 1 | >1000 |
| *Galns* | 12 | 148 |
| *Ctbs* | 3 | 135 |
| *Aqp11* | 2 | 109 |
| *Lmf1* | 6 | 309 |
| *Mgat5b* | 11 | 152 |
| *Glb1* | 15 | 377 |
| *Hs3st3a1* | 1 | >3000 |
| *Kera* | 1 | 77 |
| *Sulf2* | 17 | 46 |
| *Stogalnac3* | 1 | >3000 |
| *Bgn* | 4 | 19 |

| | | |
|---|---|---|
| *Pcsk6* | 20 | 146 |
| *Arsb* | 7 | 51 |
| *Dcn* | 7 | 198 |
| *Itih5* | 2 | 947 |

**Supplemental Table S3.** Number of Premature Termination Codons (PTCs) in the retained introns of matrix transcripts of *Sfpq* knockdown group.

| Gene | Intron Retention # | Number of Premature Termination Codon (PTC) |
|---|---|---|
| *Col4a2* | 4 | >1000 |
| *Adam12* | 1 | >1000 |
| *Ntn1* | 2 | 476 |
| *App* | 13 | 248 |
| *St6galnac3* | 1 | >3000 |
| *Galnt11* | 1 | >1000 |
| *Galnt10* | 7 | 219 |
| *Asph* | 4 | 150 |

**Supplemental Table S4:** Intron retention profiling of SFPQ, EWSR1, TAF15, and TRA2B knockdown in ST2 cells. Excel file attached.

**Supplemental Table S5:** GO analysis of genes with increased intron retention in SFPQ, EWSR1, TAF15, or TRA2B depleted ST2 cells. Excel file attached.

**Supplemental Table S6**: rMATS-based identification of alternative splicing alterations following SFPQ, EWSR1, TAF15, or TRA2B knockdown in ST2 cells. Excel file attached.

**Supplemental Table S7:** RNA binding proteins (RBP) with significantly differential binding in splicing events altered by PRMT1 (p<0.00001). Excel file attached.

**Supplemental Table S8:** Significant RBP with top occurrences and their corresponding p-values.

| | **Up in siPRMT1** | **Down in siPRMT1** |
|---|---|---|
| **Significant RBP** | SFPQ | SFPQ |
| **RI** | NA | NA |
| **SE** | 6(smallest_p_in_downstreamExonIntron)/342 (pVal= 0.0001) | 39(smallest_p_in_upstreamExonIntron)/314 (pVal= 0.00088) |
| **MXE** | 103(R10)/347 (pVal=0.0046) | 123(R2)/342 (pVal=0.006) |
| **A5SS** | 64(R5)/192 (pVal=0.006) | 60(R1)/127 (pVal=0.015) |
| **A3SS** | NA | 24(R3)/101 (pVal=0.004) |

**Supplemental Table S9:** List of primers. Excel file attached.
